# Ecological diversification in rapidly evolving populations

**DOI:** 10.1101/2025.09.15.676408

**Authors:** Daniel P.G.H. Wong, Benjamin H. Good

## Abstract

Microbial populations have an enormous capacity for rapid evolutionary change. Some mutations increase the fitness of their lineage and compete with each other in a process known as clonal interference. Other mutations can evade competitive exclusion by diversifying into distinct ecological niches. Both processes are frequently observed in natural and experimental settings, yet little is known about how they interact in the parameter regimes most relevant for microbes. Here we address this gap by analyzing the dynamics of ecological diversification in a simple class of resource competition models, where individuals acquire mutations that alter their resource uptake rates. We focus on large adapting populations, where mutations occur so frequently that their ecological and evolutionary timescales overlap. In this regime, we show that the competition between linked mutations causes the population to self-organize into a smaller number of distinct ecotypes, driven by an emergent priority effect that favors the resident strains. We demonstrate that these priority effects bias the long-term metabolic structure of the population, producing qualitative departures from existing ecological theory. We argue that similar dynamics should arise for other rapidly evolving ecosystems, where adaptive mutations accumulate at many linked genetic loci.

## Introduction

Microbial populations can adapt to their local environment by rapidly acquiring new mutations. Many such mutations are subject to competitive exclusion, and will either take over the population or go extinct. However, some mutations can evade competitive exclusion by expanding into a different ecological niche. Striking examples of this behavior have been observed in microbial evolution experiments, where initially clonal populations spontaneously diversify into multiple coexisting ecotypes (1–12). The mechanism of coexistence can often be traced to subtle differences in resource use (7, 13), but similar effects can also arise through differences in predation (14–16), antagonism (16–18), and immunological memory (19, 20). While these ecotypes coexist with each other in the short-term, their relative balance can shift over longer evolutionary timescales as their descendants acquire further mutations. Evidence for such quasi-stable coexistence has been observed in a variety of natural (21–27) and experimental (11, 12, 28–32) settings. However, our theoretical understanding of this diversification process – and its impact on microbial evolution – remain poorly understood in comparison.

Most existing approaches for modeling ecological diversification assume that evolution is much slower than ecology (33–40). Mutations arise one-by-one, and their fates are determined by an ecological competition against the other resident strains in the population. While this weak-mutation limit has shed light on the conditions where ecological diversification is likely to occur, it is known to break down in microbial populations where ecotypes are commonly observed. In many cases of interest, from laboratory evolution experiments (28, 41, 42) to natural populations of viruses (43–45), bacteria (24, 46) and certain cancers (47), the size of the population is large enough that multiple new mutations will arise and compete with each other at the same time. When recombination is limited, the competition between these linked mutations (“clonal interference”; 48) ensures that the ecological fate of a new mutation will rarely be decided before the next mutation occurs. This leads to a more complex set of dynamics, in which the ecological and evolutionary timescales are inherently intertwined.

Clonal interference has been extensively studied in the absence of ecological interactions (48–56), but much less is understood in the more general case where ecotype structure can emerge. Several recent studies have started to explore these effects using computer simulations (18, 34, 57, 58). However, our limited theoretical understanding of these overlapping timescales leaves many basic questions unresolved: Does clonal interference tend to promote ecological diversification by producing a wider range of new mutations? Or does it constrain coexistence by allowing the resident strains to constantly encroach on each other’s niches? What signatures do these dynamics leave in genomic data? And how do they impact the ecological or metabolic phenotypes that emerge in the population in the long run?

Here we address these questions by developing a population genetic theory of ecological diversification that is valid for rapidly evolving populations. We focus on a simple class of resource competition models, where cells compete for different metabolites in a well-mixed environment, while acquiring heritable mutations in their genomes. Focusing on the two resource case, we show that rapidly evolving populations naturally cluster into a smaller number of distinct ecotypes, even when their underlying strain diversity is very large. By extending traveling wave models of clonal interference, we show that this self-organization is driven by an emergent priority effect that hinders ecological diversification by favoring the resident ecotypes. We show how these priority effects bias the long-term ecological structure of the community, and discuss the implications of these results in the context of recent microbial evolution experiments. Together, these results provide a quantitative framework for modeling ecological diversification in settings where ecology and evolution are both widespread.

## Results

### Modeling ecological diversification in rapidly evolving populations

To understand how rapid evolution influences ecological diversification, we focus on a simple class of resource competition models that have been studied in a number of previous works (34– 37, 59–65). These models aim to capture several key empirical features of microbial evolution experiments (3, 5, 28), while remaining as analytically tractable as possible. In the simplest version of the model, we assume that individuals compete for ℛ substitutable resources in a chemostat-like environment (Fig. 1A). Each resource is externally supplied at a constant rate *β*_*i*_, which represents the fraction of available biomass that is supplied through resource *i*. Each strain *μ* in the population is then characterized by a resource-utilization vector 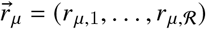, which describes how rapidly it consumes each of the available resources. Following previous work (34, 59), it will be useful to decompose this phenotype into an overall magnitude (or “budget”) *X*_*μ*≡_ log ∑ _*i*_ *r*_*μ,i*_, and a normalized “strategy” 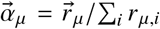, which reflects the relative effort that strain *μ* devotes to importing each of the available resources.

**Fig 1.**
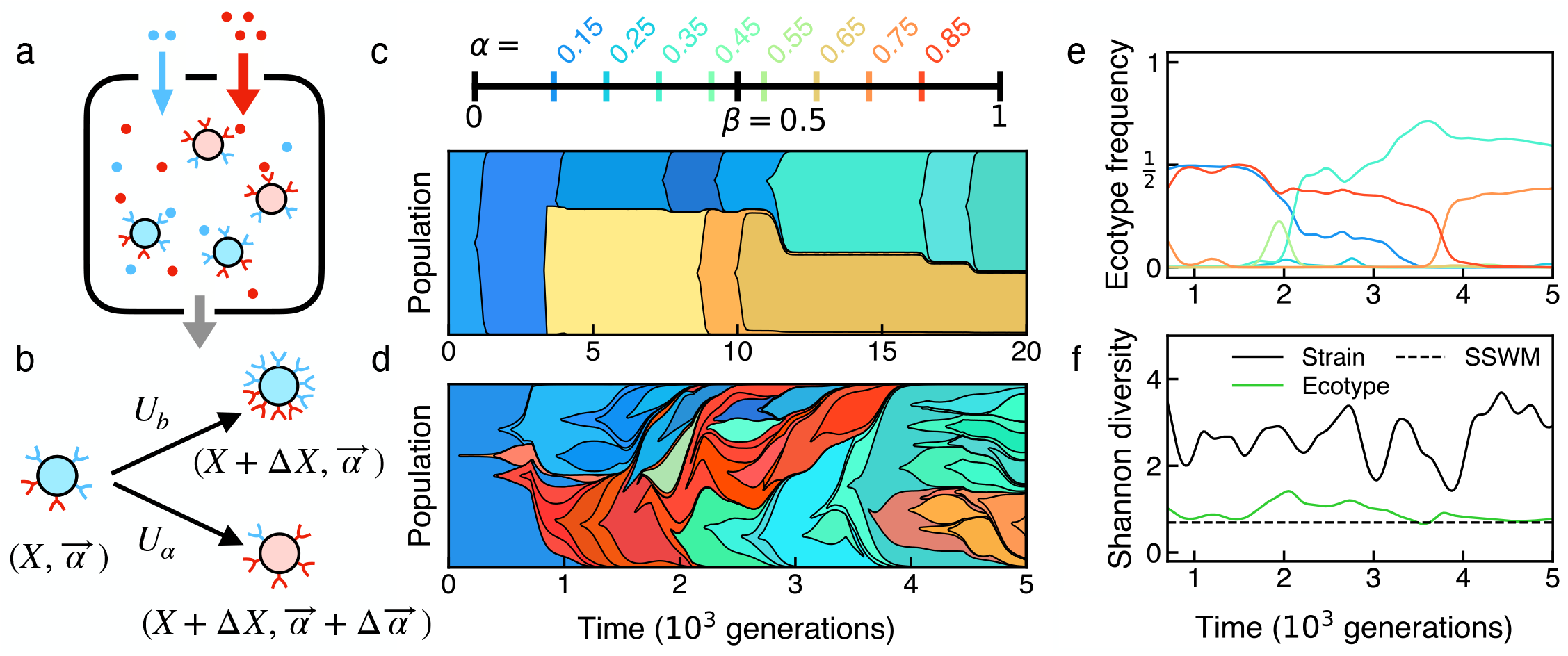
Modeling ecological diversification in rapidly evolving asexual populations. (**a**) Individuals compete for ℛ substitutable resources in a chemostat-like environment. (**b**) Individuals acquire mutations that alter their resource uptake rates 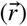. Mutations that impact the normalized resource uptake strategy 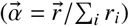 occur at rate *U*_*a*_, while mutations that only increase the overall magnitude *X* = log ∑_*i*_ *r*_*i*_ occur at rate *U*_*b*_. (**c, d**) Simulated dynamics for ℛ = 2 resources at small (c, *N* = 10^6^) and large (d, *N* = 10^9^) population sizes, where strategy mutations are sampled uniformly from the discrete strategies (or “ecotypes”) depicted at top. Muller plots show the relative abundances of each strain that reaches at least 5% frequency in the population. Strains are colored by their resource uptake strategies (top), with shading denoting different genotypes. Other parameters are *U*_*b*_ = 10^−8^, *s*_*b*_ = 0.02, *U*_*a*_ = 10^−7^. (**e**) Total relative abundances of the ecotypes in panel d. (**f**) Shannon diversity (base e) of the individual strains in panel d (black) as well as the coarse-grained ecotypes in panel e (green). The dashed line depicts the maximum stable diversity possible in the rare mutation regime in panel c, when two coexisting strains are at equal frequencies. These data show that large populations cluster into a few coarse-grained ecotypes, even when their underlying strain diversity exceeds the competitive exclusion bound.

When resource uptake is fast compared to dilution, the population rapidly reaches a fixed carrying capacity *N*, and the state of the ecosystem can be described by the collection of relative strain frequencies *f*_*μ*_. Under suitable assumptions (SI Section 1), these strain frequencies can be approximated by the stochastic differential equation,

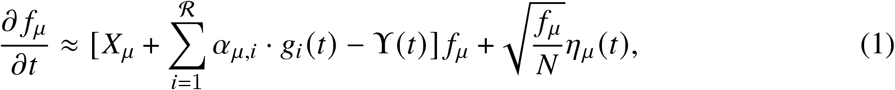

where *η*_*μ*_ (*t*) is a Brownian noise term (66) that captures the stochastic effects of genetic drift, *g*_*i*_ (*t*) ≈ (*β*_*i*_ − ∑ _*μ*_ *α*_*μ,i*_ *f*_*μ*_)/*β*_*i*_ is an intensive variable that measures the local availability of resource *i*, and *ϒ*(*t*) ≈ ∑_*μ*_ *X*_*μ*_ *f*_*μ*_ + ∑_*μ,i*_ *α*_*μ,i*_*g*_*i*_ *f*_*μ*_ is an additional gauge term that ensures that the relative frequencies remain normalized. These Lotka-Volterra-like dynamics allow for stable coexistence of up to ℛ competing strains that partition the resources in different ways (34, 59, 67).

To incorporate evolution, we assume that the individuals in the population can also acquire mutations that alter their resource consumption phenotypes. As above, it will be convenient to classify these mutations by their effects on the overall magnitude (*X*_*μ*_) and resource uptake strategy 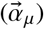 of the mutant strain (Fig. 1B). For most of this work, we will consider two broad classes of mutations. Strategy mutations, which occur at a total per capita rate *U*_*α*_, alter the resource uptake strategy of an individual and potentially its overall magnitude as well. We also allow for a second class of “constitutively beneficial” mutations, which arise at rate *U*_*b*_ and increase the total magnitude of 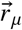 by a characteristic amount Δ*X*≈ *s*_*b*_, while leaving the relative rates intact 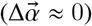. This second class of mutations models changes to cellular functions that are only tangentially related to resource consumption (e.g. eliminating a pathway that is not necessary in the current environment). Such mutations are routinely observed in microbial evolution experiments (28, 41, 68), and have previously been shown to have large impacts on the emergent eco-evolutionary dynamics (34). While the assumptions of a fixed effect size (Δ*X* ≈ *s*_*b*_) and a perfect tradeoff 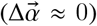 may initially seem to be artificial, we will show below that both assumptions can be relaxed without influencing our main results.

Together, these assumptions define a simple evolutionary model, similar to Refs. (34, 58), which allows us to explore the dynamics of ecological diversification across a range of population sizes and mutation rates. The simulations in Fig. 1C,D show an example of this diversification process for ℛ= 2 resources and two different simulated population sizes: a smaller population where mutations occur infrequently (*NU ≈* 0.1; Fig. 1C), and a larger population where multiple new mutations are produced in each generation (*NU* ≈ 100; Fig. 1D). These additional mutations lead to qualitatively different behavior, which we explore in more detail below.

### Rapidly evolving populations cluster into a few distinct ecotypes

In small populations (*NU* ;S 1), the diversification dynamics in Fig. 1C are well-described by the strong-selection, weak-mutation (SSWM) framework in Ref. (34). The salient features are present even for ℛ = 2 resources, which we focus on for most of this work. In this case, both the resource supply rates and uptake vectors can be characterized by scalar values, *β* and *α*_*μ*_ (Fig. 1C, top) corresponding to the first resource dimension, with the remaining components fixed from normalization. When mutations occur infrequently, the population reaches its stable equilibrium long before the next mutation occurs. This separation of timescales leads to the punctuated dynamics in Fig. 1C, where extended periods of stasis are interspersed with rapid switches in composition. When a new mutation arises in such a setting, its initial growth rate (or invasion fitness) is given by

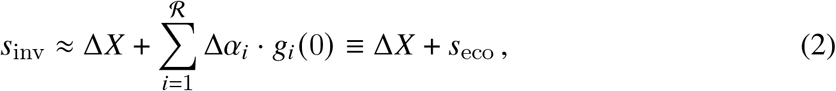

where Δ*X* = *X*_2_ − *X*_1_ and 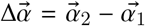 denote the phenotypic effects of the mutation, and the resource availabilities *g*_*i*_ (0) are set by the previous ecological equilibrium (34). The first term in this expression acts like a constitutive cost or benefit, while the ecological component (*s*_eco_ ≡ ∑_*i*_ Δ*α*_*i*_*g*_*i*_) is a variable quantity that depends on the current resource utilization of the population.

Most such mutations will drift to extinction even when they are deterministically favored to invade (*s*_inv_ *>* 0). However, a lucky fraction (*p*_est_ ≈ 2*s*_inv_) will establish in the population and eventually grow to larger frequencies, producing a new ecological equilibrium. The properties of this state will strongly depend on the phenotype of the invading strain (34, 35, 60). Some mutants will outcompete the current residents and fix (*f* ^*^ ≈ 1), while others will stably coexist with one or more of the residents by altering the resource availabilities in Eq. (1). The outcomes are particularly simple for ℛ = 2 resources, since at most two strains can coexist at a time. The equilibrium frequency is given by

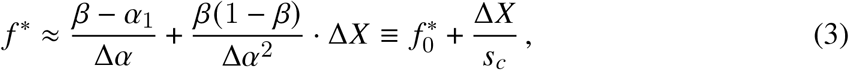

with coexistence occurring whenever *f* ^*^ *<* 1 (34; SI Section 2). This leads to a new set of *g*_*i*_ values, which can be substituted into Eq. (2) when the next mutation arises.

Equation (3) also defines two new parameters that will be important for much of our analysis below. The first is the baseline equilibrium frequency, 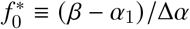, that is attained when the two strains share the same total fitness (Δ*X* ≈ 0). The second is a characteristic fitness scale, *s*_*c*_ ≡Δ*α*^2^ /*β* (1− *β*), which describes how constitutive fitness differences perturb this baseline equilibrium. This susceptibility parameter controls how the balance between two strains shifts when they acquire a constitutively beneficial mutation (Δ*X*→ Δ*X* ±*s*_*b*_), as well as the total fitness differences that are tolerated before one of the strains is driven to extinction (Fig. 1C).

While this existing theory applies to Fig. 1C, qualitatively different behavior emerges in larger populations when mutations occur more frequently (*NU* ≳ 1; Fig. 1D). The most striking difference is the absence of any apparent ecological equilibria: at any given timepoint, all of the resident strains exhibit nonzero relative growth rates, signaling that ecology and evolution are now operating on similar timescales. A consequence of this behavior is that the local strain diversity often exceeds the competitive exclusion bound set by the number of available resources (Fig. 1F), for the simple reason that new mutations are introduced into the population much faster than they can be purged (57–59).

Despite this high strain diversity, we find that when individuals are grouped together by their resource strategies 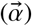, the population is usually dominated by only a few distinct strategies at a time (Fig. 1E,F), even when strategy mutations are common (*NU*_*α*_ ≫ 1). Moreover, these dominant strategies can persist at high frequencies for thousands of generations (Fig. 1E), while their underlying strains turn over at a much faster rate (Fig. 1D). This emergent simplicity highlights the utility of working in the resource strategy basis. It also suggests that further progress can be made by tracking the emergence and maintenance of a smaller number of distinct “ecotypes”, which share the same resource uptake strategies but contain strains with many different genotypes and fitnesses (*X*_*μ*_).

### A dynamical priority effect hinders the invasion of new ecotypes

To understand the mechanisms driving the behavior in Fig. 1C-F, we start by considering the simplest case, where a new strategy mutation arises in a population with only a single resident ecotype. We initially assume that further strategy mutations can be neglected (*NU*_*α*_ → 0), so the fate of the new ecotype is shaped by fitness mutations alone. These “pairwise” invasions will provide the building blocks necessary for understanding the more general scenarios below. In large populations (*NU*_*b*_ ≫1), the initial stages of diversification are best understood by considering the population’s distribution across the strategy (*α*) and fitness (*X*) dimensions of phenotype space (Fig. 2A; SI Section 3). When the new ecotype first arises, previous work has shown that the resident ecotype will have formed a “traveling fitness wave” along the *X* axis, with a characteristic shape *ρ* (*x*) that moves to the right a steady state rate *v* (50–54). This distribution has a maximum width *x*_*c*_, which roughly coincides with the location of the fittest individuals in the population: these individuals in the “nose” of the fitness distribution possess *q* ≈*x*_*c*_ /*s*_*b*_ more beneficial mutations than the population average, and will grow to become the dominant fraction of the population in a characteristic “sweep time” *T*_sw_ ≈*x*_*c*_ /*v* (Fig. 2A, left; SI Section 3.1). A new ecotype will arise on one of these extant genetic backgrounds, with a relative fitness drawn from *ρ* (*x*). It will then go on to found its own fitness wave with resource strategy *α*_2_ = *α*_1_ Δ*α* (Fig. 2A, right).

**Fig 2.**
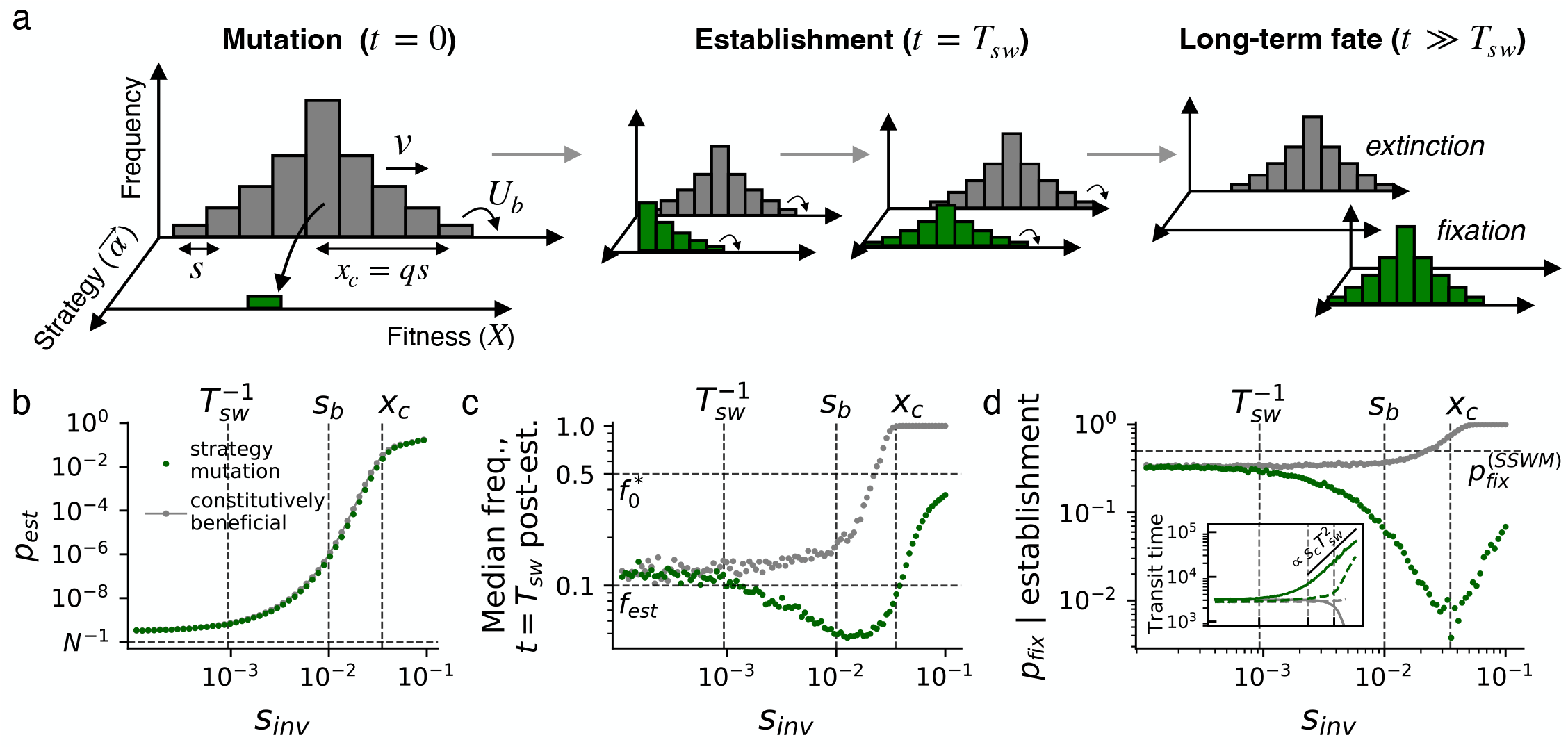
Clonal interference creates a priority effect that favors the resident ecotype. (**a**) Schematic of a strategy mutation arising in a population with a single resident ecotype. The mutation arises in an individual from the resident fitness distribution (grey), which has a maximum width *x*_*c*_ and moves to the right at rate *v*. Its descendants found a new ecotype (green) that competes with the resident ecotype as they both acquire additional mutations. (**b**) The probability that a strategy mutation “establishes” (i.e. reaches 10% frequency) as a function of its initial invasion fitness *s*_inv_. Green points show simulation results for *N* = 10^10^, *U*_*b*_ = 10^−6^, and *s*_*b*_ = 0.01; for each value of *s*_inv_, the values of *α*_1_, *α*_2_, and *β* were chosen so that *f* ^*^ 0 = 1/ 2. Grey points show the establishment probabilities of constitutively beneficial mutations with the same values of *s*_inv_. (**c**) The median frequencies of the mutations in panel b *T* sw ≈ 1000 generations after first reaching 10% frequency. The dashed line denotes the expected frequency in the absence of clonal interference. (**d**) The long-term fixation probabilities of the mutations in panel c. Inset: the average length of time required for a mutation to transit from 10% frequency to either fixation (solid curves) or extinction (dashed).

When the frequency of the new ecotype is still small, the growth rate of its founding strain can be calculated from a generalization of Eq. (2),

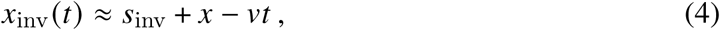

with *s*_inv_ ≈ Δ*X* +*s*_eco_ ≈ Δ*X* + Δ*α* (*β* − *α*_1_) / (*β* 1− *β*). The two new terms in Eq. (4) account for (i) the initial relative fitness of the mutant’s previous genetic background, and (ii) the ongoing adaptation of the resident ecotype, which causes the relative fitness of any given strain to systematically decline with time. This declining growth rate implies that many initially adaptive mutants that survive genetic drift will eventually be outcompeted by the resident population before they reach appreciable frequencies (“clonal interference”). To reach a substantial fraction of the population, a new ecotype must either arise on such an anomalously fit genetic background that it “outruns” the resident population, or it must quickly acquire one or more additional beneficial mutations that accomplish the same effect. As we show below, these chance events all occur while the mutant lineage is still at a low frequency. This implies that both fitness and strategy mutations — and any other lineage with the same value of *x*_inv_ (0) — will have the same probability of establishing.

We verify this prediction in Fig. 2B, by comparing the observed establishment probability of a new strategy mutation — operationally defined as the probability that it reaches 10% frequency — against a constitutively beneficial mutation with the same value of *s*_inv_. For concreteness, we consider a simple class of strategy mutations with Δ*X* = 0 and Δ*α* = 2 (*β* −*α*_1_), such that *s*_inv_ ≈*s*_eco_ ≈ Δ*α*^2^ / 2*β* (1 − *β*). Fig. 2B shows that the establishment probabilities of these mutations closely track their non-ecological counterparts, and exhibit the classic hallmarks of clonal interference (53, 69): the establishment probability approaches the neutral limit (*p*_est_ ∼1 /*N*) when *s*_inv_ ≲ 1 / *T*_sw_, and then increases sharply at higher values of *s*_inv_ before returning to the SSWM expectation (*p*_est_ ≈ 2*s*_inv_) when *s*_inv_ ≳ *x*_*c*_. This emergent neutrality at small values of *s*_inv_ implies that sufficiently small changes in resource strategy [Δ*α* ;S *β* (1 − *β*) / (*β* −*α*_1_) *T*_sw_] will be effectively invisible to natural selection, even when their ecological selection pressures are nominally large (*Ns*_eco_ ≫1).

At higher frequencies, established strategy mutations start to diverge from their non-ecological counterparts, as environmental feedbacks from the invading ecotype eventually reduce the value of *s*_eco_ (*t*) ≡ ∑_*i*_ Δ*α*_*i*_*g*_*i*_ (*t*). In the weak mutation regime, these feedbacks would normally cause the new ecotype to stabilize at the equilibrium frequency in Eq. (3), which is equal to *f* ^*^ ≈ 50% for all of the mutations that are simulated in Fig. 2B. Interestingly, however, Fig. 2C shows that the long-term frequency of the invading ecotype is consistently smaller than 50%, and reaches its lowest point for intermediate selection strengths where *s*_inv_ ≈ *x*_*c*_. On even longer timescales (*t ≫T*_sw_), thelower initial frequency of the invading ecotype eventually impacts its ability to accumulate of future mutations. This implies that the invading ecotype is much more likely to be driven to extinction by the resident in the long run (Fig. 2D). We call this emergent bias in favor of the resident ecotype a priority effect (70), since it does not arise from any intrinsic differences between the two ecotypes, but rather from their order of arrival in the population.

The origin of this priority effect can be understood by considering the genetic backgrounds of the invading ecotypes. The total establishment probability of a new mutation can be expressed as an average over genetic backgrounds,

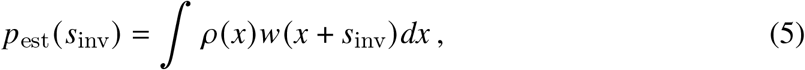

where *w* (*x*) represents the conditional establishment probability of a lineage with *x*_inv_ (0) = *x* (SI Section 3.3). Previous work (53) has shown that when *s*_inv_ ≲ *x*_*c*_, the integral in Eq. (5) is strongly peaked around a characteristic value *x*^*^ ≈ *x*_*c*_ −*s*_inv_, such that the initial growth rate of a successful mutation is likewise around 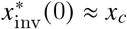. In other words, successful mutations appear to “tune” their genetic background so that their net growth rate coincides with the “nose” of *ρ* (*x*), where the fittest individuals in the population currently reside.

When projected onto the *α* and *X* axes in Fig. 2A, this tuning behavior implies that the *X* coordinate of a successful strategy mutation will be strongly peaked around *X*^*^ ≈ *x*_*c*_ − *s*_inv_, which lies below the nose of the resident ecotype by a characteristic amount *s*_inv_. This constitutive fitness deficit only becomes important *T*_sw_ ≈ *x*_*c*_ /*v* generations later, when both sets of strains have grown to occupy the dominant fraction of the population. At this point, the invading ecotype experiences an effective fitness difference of Δ*X*_eff_ ≈ −*s*_inv_ compared to its competitors, which would lead to a lower equilibrium frequency according to Eq. (3). This heuristic argument helps explain the emergent priority effect observed above: successful strategy mutations tend to “mortgage” their ecological advantage to maximize their short-term probability of establishing. But they eventually pay a price at higher frequencies, when their rapid expansion causes their ecological advantage to suddenly dissipate.

### Oscillatory invasions of new ecotypes

While this “fitness mortgaging” picture is essentially correct, quantitatively predicting the behavior in Fig. 2C,D requires a more careful analysis of an ecotype’s invasion dynamics over time. In rapidly evolving popluations, the frequency of a successful ecotype obeys a time-dependent generalization of Eq. (3),

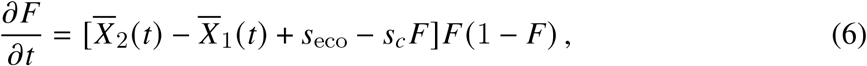

where 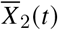 and 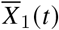 denote the average *X* coordinates of the invading and resident ecotypes, respectively (SI Section 3). Fig. 3 shows that these frequency trajectories can undergo large oscillations when the mutant ecotype first invades, reaching a peak frequency around *T*_sw_ generations, before declining to a quasi-steady state level at later times. We will now show how these oscillatory dynamics emerge from a delayed feedback between two distinct subsets of the population: (i) the ecologically-relevant bulk, which shapes the current environment, and (ii) the evolutionarily-relevant nose, which will come to dominate the population at later times.

**Fig 3.**
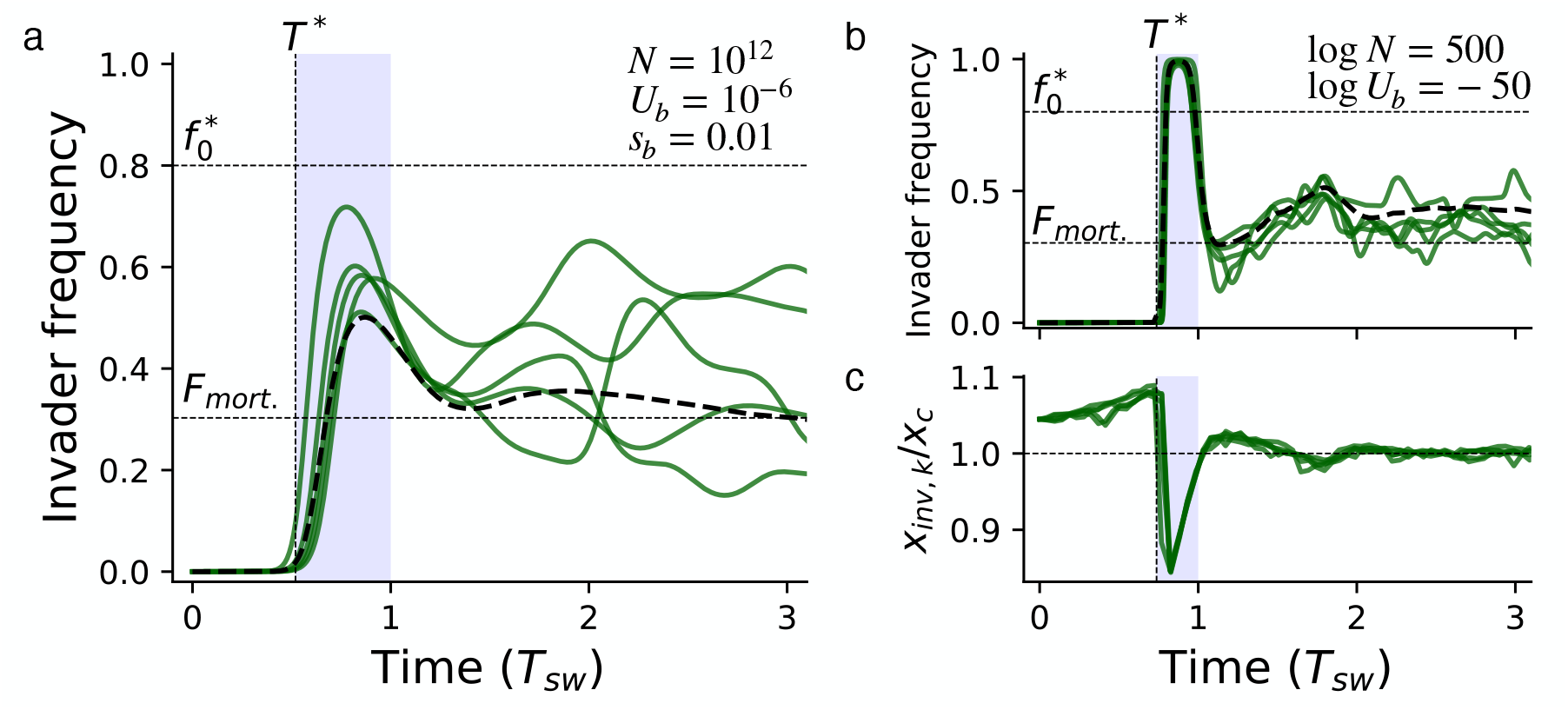
Oscillatory invasion dynamics of new ecotypes. (**a**) Replicate frequency trajectories of an invading ecotype with *s*_eco_ ≈ 0.03 and 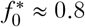 that arises on an initial genetic background with *x*_inv_ (0) ≈ *x*_*c*_ + *s*_*b*_. Simulated parameters are *N* = 10^12^, *U*_*b*_ = 10^−6^, and *s*_*b*_ = 0.01, for which *x*_*c*_ ≈ 4*s*_*b*_. Five random realizations are shown in solid lines, while the dashed line depicts an average over 50 independent replicates, which emphasizes the typical overshooting behavior between *T* ^*^ (Eq. 7) and *T*_sw_ ≈*x*_*c*_ /*v*. Horizontal dashed lines show the original equilibrium frequency *f* ^*^ (Eq. 3) and the “fitness mortgaging” term from Eq. (9). (**b**) The oscillations become more pronounced in the asymptotic limit, where log *N* = 500, log *U*_*b*_ = −50, and *x*_*c*_ ≈ 21*s*_*b*_. (**c**) The initial growth rates of successive fitness classes (Fig. S1, SI Section 3.3) for the ecotype trajectories in panel b, illustrating how the runaway growth is eventually stabilized ∼*T*_sw_ generations later.

We quantify these dynamics by considering the trajectories of successive fitness classes, *f*_*k*_ (*t*), which denote the collection of mutant individuals that have acquired *k* additional mutations. We focus on the limiting behavior of these cohorts in the empirically-relevant scenario where *s*_*b*_ *≫ U*_*b*_ *≫* 1 *N*, where the key fitness scales in Fig. 2A satisfy 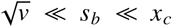 (52). In this regime, the fitness of the invading ecotype is usually dominated by a single value of *k*, corresponding to the largest fitness class at that instant in time. We can combine this approximation with Eq. (6) to obtain a piecewise description of the ecotype’s frequency trajectory over time (SI Section 3.4).

In the initial phase of each trajectory (*t < T*_sw_), the invading ecotype is dominated by the founding *k* = 0 class, and the first three terms in Eq. (6) are well-approximated by the initial growth rate *x*_inv_ (*t*) in Eq. (4). The resulting dynamics strongly depend on how *x*_inv_ (0) compares to *x*_*c*_. When *x*_inv_ (0) *> x*_*c*_, the invading ecotype will eventually reach a frequency where the ecological feedbacks become important. Before this point, the invading ecotype is well-approximated by the linear model in Eq. (4), and increases exponentially with a steadily declining growth rate *x*_inv_(*t*). After a critical time

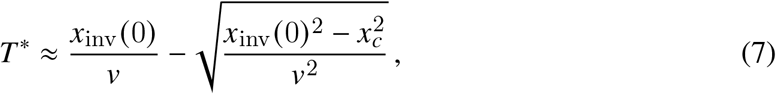

the ecological feedbacks in Eq. (6) lead to a rapid equilibration of the ecotype frequency around a quasi-steady-state value,

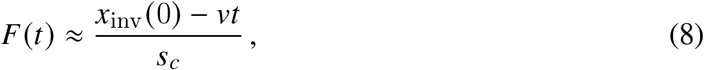

which applies when 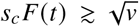.The decreasing form of this expression implies that the invading ecotype will attain a maximum value 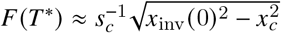 and will then begin to decline over time, as the resident ecotype continues to adapt relative to the initial founding clone.

On longer timescales, the *k* = 0 class will eventually be replaced by mutant lineages that have acquired additional beneficial mutations. In our parameter regime of interest (*s*_*b*_ ≫*U*_*b*_ 1≫ / *N*), these replacement dynamics take on a particularly simple form: the *k*th class starts to dominate the new ecotype at a characteristic time *t* ≈*T*_sw_ + *T*_*k*_, where *T*_*k*_ denotes the corresponding establishment time where *f*_*k*_ (*T*_*k*_) ≈1/ 2*Nx*_inv,k_ (*T*_*k*_) (Fig. S1; SI Section 3.4). After the *k*th class takes over, the fitness of the invading ecotype will have advanced by a total amount *ks*_*b*_, while the fitness of the resident ecotype will have increased by *v* (*T*_sw_ +*T*_*k*_)over the same time interval. This leads to a modified version of Eq. (8) that applies for *t* ≳ *T*_sw_:

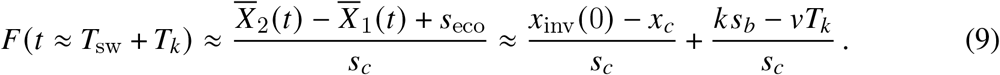

This expression constitutes the rapid-evolution analogue of the ecological equilibrium in Eq. (3). The first term matches the naive expectation from our “fitness mortgaging” argument above, while the second term accounts for differences in the rates of establishment between the invading and resident ecotypes.

The statistics of the first few *T*_*k*_ values can be calculated by generalizing the calculations in Ref. (52) (SI Section 3.3). The *k* = 1 class establishes slightly earlier than the resident nose, since its feeding class grows at a higher initial rate (*x*_inv_ (0) *> x*_*c*_; Fig. 3C). This difference in establishment times causes the invading ecotype frequency in Eq. (9) to be slightly larger than expected from the fitness mortgaging term alone. These early establishments continue for several additional fitness classes, as each successive establishment widens the gap in the initial growth rate (Fig. 3C), leading to an increase in the overall frequency. However, the trend eventually reverses for *T*_*k*_ ≳ *T* ^*^, when the ecological feedbacks cause the effective growth rate to drop to *ks*_*b*_ *< x*_*c*_ (SI Section 3.4). These damped oscillations can persist for several additional multiples of *T*_sw_, as observed in Fig. 3. However, the corrections are often modest in practice, such that the first term in Eq. (9) provides a reasonable approximation to the ecotype frequency when *t* ∝ *T*_sw_.

These results show that the frequency trajectory of the invading ecotype strongly depends on the critical quantity *δ* (*s*_inv_) ≡*x*_inv_ (0) − *x*_*c*_, which represents the growth advantage of mutant over the resident nose, and implicitly depends on *s*_inv_ through the distribution of successful background fitnesses *x* in Eq. (5). A more careful analysis in SI Sections 3.2 and 3.3 reveals that the distribution of positive *δ* values can be approximated by a truncated Gaussian distribution,

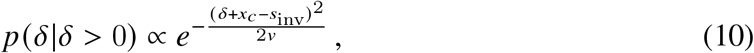

which is peaked near *δ* ≈*s*_inv_ −*x*_*c*_ for *s*_inv_ <‘*x*_*c*_ and *δ* ∼*v* /*x*_*c*_ otherwise. These results imply that mutants with larger values of *s*_*c*_ ≡ Δ*α*^2^ /*β* (1− *β*) must sample proportionally larger values of *δ* to reach a given post-invasion frequency *F* (*t*). The fitness mortgaging effect in Eq. (10) makes this increasingly unlikely until *s*_inv_ ≳ *x*_*c*_, which explains the non-monotonic behavior in Fig. 2C. At the same time, Eq. (7) shows that populations with stronger clonal interference (*x*_*c*_ ≫ *s*_*b*_) can amplify smaller values of *δ* to equivalent peak frequencies *F* (*T* ^*^) during the initial transient phase of the invasion. This highlights the counter-intuitive dynamical effects that emerge in rapidly evolving populations.

### Long-term dynamics of coexisting ecotypes

On longer timescales (*t* ≫*T*_sw_), the cumulative differences in the rates of establishment will start to dominate the local equilibrium in Eq. (8), and the ecotype frequencies will gradually diverge from their initial post-invasion levels. In the absence of additional strategy mutations, these fitness differences will eventually grow so large that they will drive one of the two ecotypes to extinction (Fig. 2D).

Since these long-term differences reflect the contributions of many accumulated mutations (*k* ≫1), it becomes natural to model their dynamics as a generalized random walk. Building on previous results derived in non-ecological settings (52, 54), we show that the long-term fitness of each ecotype can be modeled as an effective diffusion process,

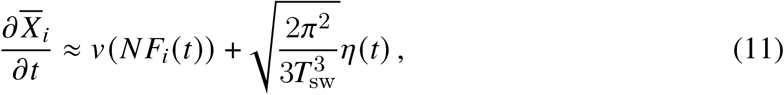

where *v* (*N*) denotes the rate of adaptation as a function of the population size *N* (SI Section 3.5). Substituting this expression into Eq. (9) yields a corresponding model for the ecotype frequency,

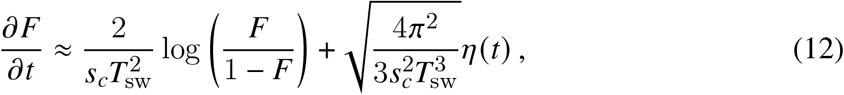

which applies for *t* ≫*T*_sw_ ≫1 /*s*_*c*_. The first term in this model reflects the deterministic difference in the rates of adaptation of the two ecotypes due to their unequal population sizes. The second term arises from the stochasticity in the timing of individual establishment events (52, 54), which are correlated on intermediate timescales (Δ*t* ∼*T*_sw_) but behave diffusively when Δ*t T*_sw_.

Analysis of Eq. (12) reveals that the deterministic bias dominates over the stochastic term when the difference in ecotype frequencies is larger than 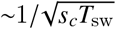 (SI Section 3.5). Beyond this critical point, the biased rates of adaptation will deterministically drive the minority strain to extinction on a timescale *T*_ext_∼*s*_*c*_*T* ^2^ ≫ *T*_sw_. The gap between these timescales justifies the original diffusion approximation in Eq. (12), and illustrates why established ecotypes can coexist for much longer than the lifetime of a typical lineage (Fig. 2D inset). At the same time, these results also show that the window of deterministic extinction becomes increasingly large when *s*_*c*_*T*_sw_ ≫ 1. This implies that even small initial frequency differences (Fig. 2C) can generate strong priority effects in the long-term composition of the population (Fig. 2D).

### Rediversification from recurrent mutations

For finite mutation rates (*NU*_*α*_ *>* 0), additional strategy mutations can arise before one of the resident ecotypes goes extinct, which alters the simple pairwise picture above. To build intuition for this more realistic case, we start by assuming that there are still just two possible ecotypes, while allowing mutations (and reversions) to occur between the two strategies at a symmetric rate *U*_*α*_ (Fig. 4A). Simulations of this model recapitulate some of the behaviors observed in the multi-ecotype setting in Fig. 1. In both cases, we observe a priority effect similar to Fig. 2, where one ecotype persists at a higher frequency for extended “epochs” of length *T*_epoch_ ≫*T*_sw_ (Fig. 4B, top), with rare transitions between successive epochs where the majority ecotypes flip. However, within the minority ecotype, there is constant genetic turnover, as new strategy mutations from the majority background continually arise and replace the previous lineages (Fig. 4B, bottom). These asymmetric cross-invasions stabilize the frequency of the minority ecotype, but lead to even stronger priority effects at the genetic level, since lineages in the minority background will have a vanishingly small probability of contributing to the future genetic composition of the population.

**Fig 4.**
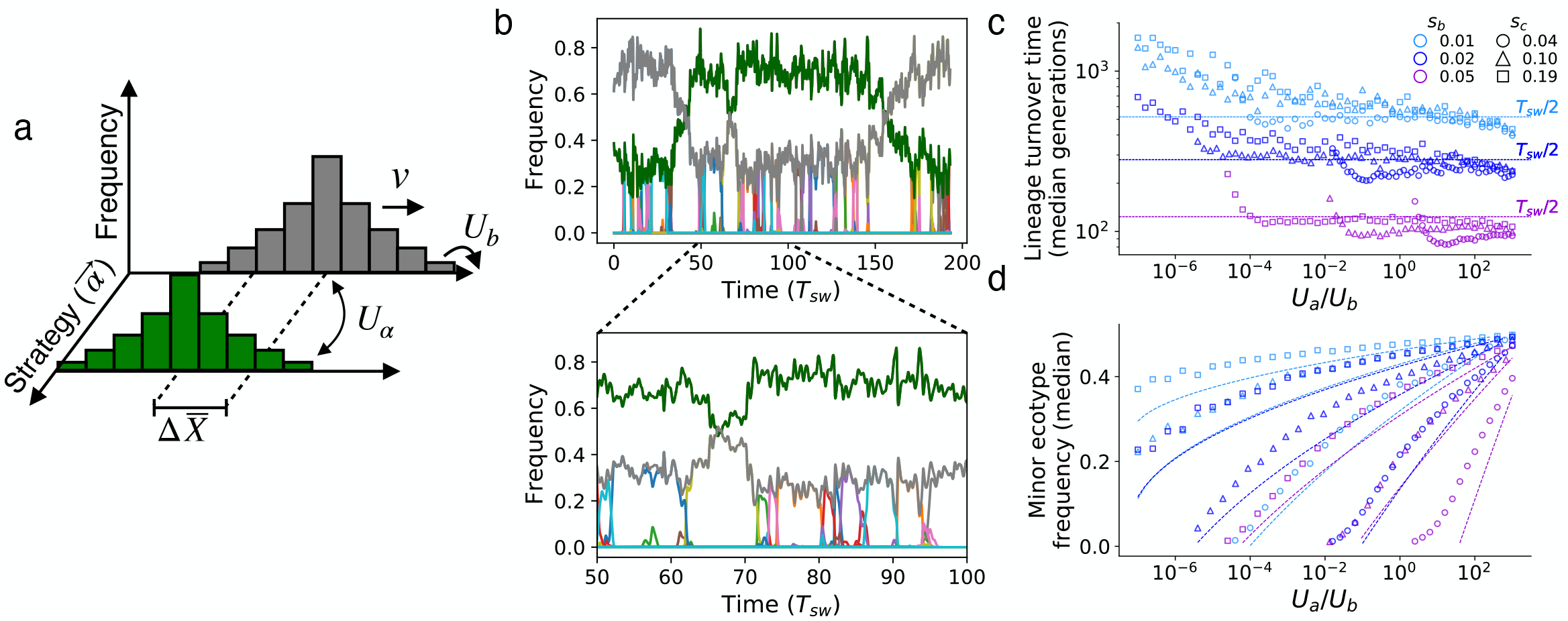
Recurrent strategy mutations stabilize coexistence by replacing the lagging ecotype. **(a)** Recurrent strategy mutations between established ecotypes can replace the resident lineage if the fitness differences grow sufficiently large. **(b)** Example simulation showing bistable ecotype frequencies on long timescales, with continuous lineage turnover on shorter timescales due to successful cross-invasions. Thick shaded curves represent overall ecotype frequencies, while thinner lines denote lineages founded by distinct strategy mutations. Ecotype parameters are *s*_*c*_ = 0.1, Δ*X* = 0, and 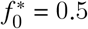, with *U*_*α*_ = 10^−11^; the remaining parameters are the same as Fig. 3A. **(c**,**d)** Lineage turnover time (**c**) and overall frequency (**d**) of the trailing ecotype for different values of *s*_*c*_ and *s*_*b*_. Symbols denote the median across simulations with the same *N, U*_*b*_ parameters as panel b. Dashed lines denote theoretical predictions (SI Section 3.6).

We can understand these new dynamics by combining several of the results derived above. In this case, a strategy mutation arising in the majority ecotype at time *t*_0_ will have an initial

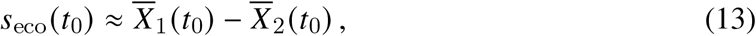

which reflects the fact that the second niche is now already occupied (SI Section 3.6). Since we have seen that the fitness difference between the two ecotypes changes on timescales much longer than *T*_sw_, the establishment probability of this mutation will still be captured by Eq. (5), with an effective invasion fitness *s*_inv_ (*t*_0_) ≈ Δ*X* +*s*_eco_ (*t*_0_). The average time to the next successful cross-invasion (*T*_cross_) can then be defined from the standard condition that approximately one mutation establishes within that interval,

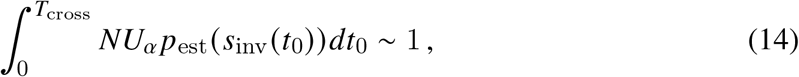

which implicitly depends on the initial fitness difference between the two ecotypes. At steady-state, the change in fitness during *T*_cross_ must balance the net fitness increase from a successful cross-invasion, which is proportional to the fitness difference *δ* (*s*_inv_) in Eq. (10). Combining this criterion with Eq. (14) yields a self-consistent solution for *T*_cross_.

These steady-state dynamics are particularly simple in the empirically-relevant regime where *NU*_*α*_ *≫*1 and *U*_*α*_ *≪s*_*b*_ (SI Section 3.6). Provided that the direct cost of the mutation is not too large, the fitness difference between the ecotypes remains approximately constant between invasions, with *s*_inv_ (*t*) *≪x*_*c*_. The self-consistency condition then reduces to *T*_cross_ ∼*T*_sw_, which is independent of the underlying resource strategies or their mutation rate *U*_*α*_ (Figs. 4C and S2). This result shows that rapid evolution pushes the ecosystem to a point where the effective fitness benefits and mutation rates of strategy mutations balance each other, so that successful cross invasions occur approximately once every *T*_sw_ generations. The emergent fitness benefits of these mutations depend only logarithmically on *U*_*α*_,

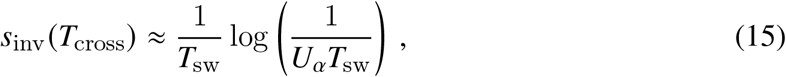

and fall in the exponential portion of *p*_est_ (*s*_inv_) in Fig. 2B. This steep dependence on *s*_inv_ implies that even small changes in the relative fitnesses of the two ecotypes will generate large shifts in *T*_cross_, which effectively stabilizes the relative fitnesses – and the corresponding ecotype frequencies in Eq. (9) – close to their average steady-state values (Figs. 4D and S2).

The weak dependence of the ecosystem on the strategy mutation rate strongly deviates from the traditional SSWM regime (34). Instead, it echoes the key lessons derived from non-ecological models of clonal interference: since most strategy mutations are wasted on insufficiently fit genetic backgrounds, ecosystem dynamics are only logarithmically dependent on the underlying mutation supply rate.

### Dynamics of multiple competing ecotypes

We can also extend the above analysis to the more general case where multiple types of strategy mutations are available. In this case, an arbitrary strategy mutation arising in the majority ecotype, Δ*α*, = *α*_3_ − *α*_1_, will satisfy a generalization of Eq. (13),

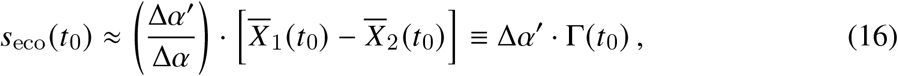

where the quantity 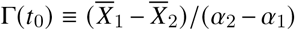 now depends on both the fitness difference and resource strategies of the resident ecotypes. These new ecotypes will continue to establish with probability *p*_est_ (Δ*X*, + Δ*α*, Γ (*t*_0_)) in Eq. (5), and will replace either the majority or minority ecotype (or both) depending on the relative values of Δ*α*, and Δ*X*,.

While a wide range of dynamics are now possible, we find that many populations reach a steady-state similar to Fig. 4, where a single majority ecotype seeds periodic cross-invasions into the opposing niche, producing a stochastic sequence of minority ecotypes. In this case, the steady-state population has a constant value of Γ (*T*_cross_), which satisfies a generalization of Eq. (14):

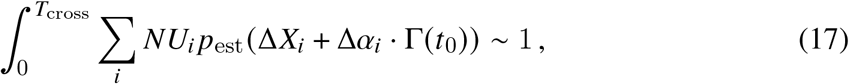

with *U*_*i*_ denoting the corresponding mutation rate for strategy mutations with effect (Δ*α*_*i*_, Δ*X*_*i*_). The presence of multiple mutation classes produces new effects that were not present in the pairwise case above. In particular, the exponential growth of *p*_est_ (*s*_inv_) implies that natural selection will strongly favor mutations with larger values of Δ*α*_*i*_, corresponding to ecotypes with more specialist metabolic strategies. This bias is much larger than the linear scaling expected in the weak mutation limit (34), and is often enough to outweigh much larger-fold differences in the underlying mutation rates (Fig. 5). As a consequence of this rapid dependence on *s*_inv_, the sum in Eq. (17) will often be dominated by a narrow range of Δ*α* and Δ*X* values, so that only a small fraction of the available ecotypes are likely to be observed at high frequencies.

**Fig 5.**
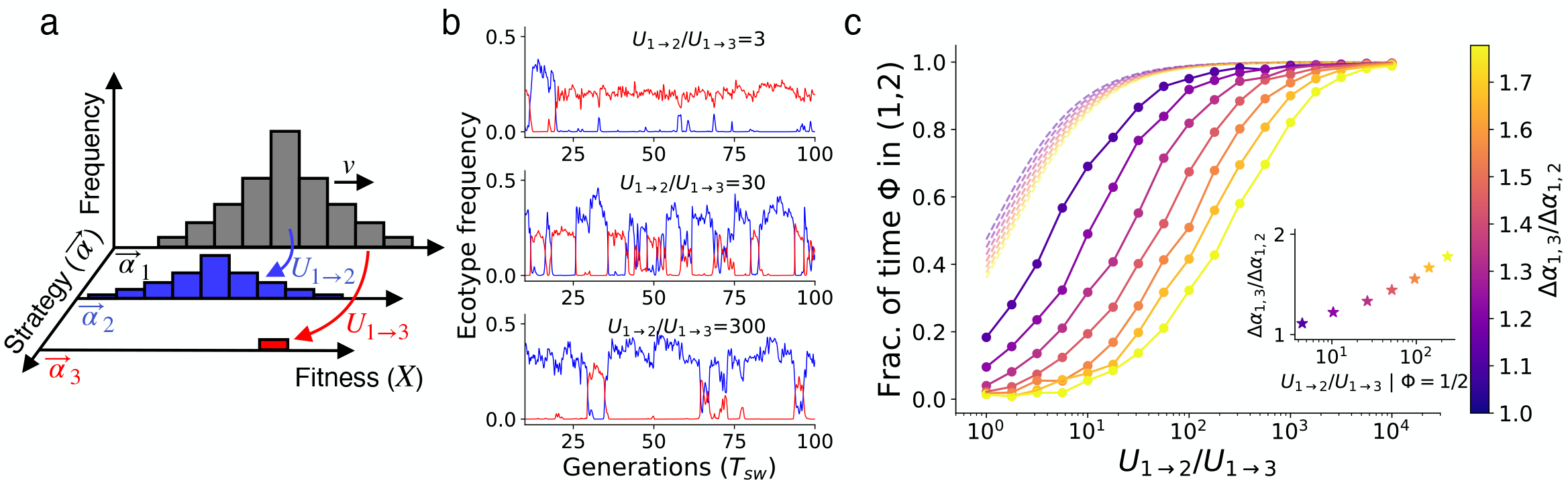
Clonal interference biases the metabolic structure of the population. (**a**) Schematic of a mutational landscape with three possible resource strategies. (**b**) Ecotype frequency trajectories from three example populations with *U*_1→ 2_ /*U*_1 → 3_ = 3, 30, and 300, with *U*_1 → 3_ = 10^−10^ held fixed. The resource strategies satisfy *α*_1_ = 0.43, *α*_2_ = 0.61, and *α*_3_ = 0.69, with *β* = 0.5, while the remaining parameters are *N* = 10^12^, *U*_*b*_ = 10^−6^, and *s*_*b*_ = 10^−2^. (**c**) The fraction of time (Φ) spent in the (*α*_1_, *α*_2_)-dominated state, for different values of *α*_3_ (color-scale) and *U*_1→2_ (x axis); all other parameters are the same as panel b. Inset: the relative mutation rate where the two lagging ecotypes are equally likely (Φ ≈1/2) for different values of U3. Clonal interference amplifies selection on small differences in resource consumption, which can overwhelm much larger differences in the relative mutation rates of different strategies.

Despite these differences, the timescale of cross-invasion remains similar to the pairwise case above. In both cases, the population self-organizes to a state where the ecological selection pressures are low enough to ensure that only ∼1 new ecotype establishes every *T*_cross_ ≳ *T*_sw_ generations, even when *NU*_*α*_ *≫* 1. Moreover, since the window for successful establishment is comparable to the *T* ^*^ ∼*T*_sw_ timescale in Fig. 3, the first such ecotype to reach high frequency will rapidly exclude its competitors, ensuring that only *R* ≈2 ecotypes will dominate the population at any given time. This creates an emergent priority effect that favors the resident ecotypes, sustaining long periods of ecotype stability in the face of rapid within-ecotype evolution (Fig. 1).

## Discussion

In large microbial populations, ecological and evolutionary dynamics often unfold on similar timescales. Our study introduces a theoretical framework for modeling these coupled dynamics in a widely used class of resource competition models, where individual cells acquire mutations that alter their resource consumption phenotypes. By focusing on the simplest case of two substitutable resources, we showed how the stochastic competition between large numbers of linked mutations impacts the emergence of new ecotypes and the structure of the resulting ecosystem.

Accounting for these realistically large mutation rates led to qualitative departures from existing models of ecological diversification (33–40), which assume that evolution is much slower than ecology. However, the direction of this effect often conflicted with the simple view that a higher mutation rate will tend to enhance ecological diversity (34, 57, 58). Instead, we found that clonal interference creates a series of “priority effects” that favor the resident lineages, and cause the population as a whole to cluster into a handful of discrete metabolic strategies (or “ecotypes”). Our analysis shows that these priority effects arise from the delayed feedback between the ecologically relevant bulk of the population – which shapes the current environment – and the high fitness “nose” that comes to dominate the population at later times. These delays combine to give the resident strains an additional opportunity to encroach on an invading mutant’s niche. Since these delayed feedbacks are a generic feature of evolution in large asexual populations (54, 71, 72), we expect that similar priority effects will likely emerge in other ecological settings (19, 73) beyond the resource competition models studied here.

Our results also provide a framework for interpreting data from microbial evolution experiments, where ecological diversification is often observed. For example, we have seen that the delayed feedbacks from clonal interference can cause invading ecotypes to undergo substantial oscillations in frequency when they first establish (Fig. 3). Similar dynamics have previously been observed experimentally, e.g. during the emergence of the crossfeeding interactions in Lenski’s long-term evolution experiment in *E. coli* (LTEE) (3, 28). Our results suggest that these dramatic reversals might be a natural consequence of ecological diversification in large populations, even in the absence of additional factors like epistasis or environmental fluctuations. On the other hand, our results also highlight cases where theory and data appear to diverge. For example, the coexisting lineages in the LTEE persist for much longer (and with larger fluctuations) than expected under our simple model, which predicts that the lagging ecotype should be replaced by cross-invaders every ∼*T*_sw_ generations. These examples illustrate how a quantitative understanding of diversification can uncover qualitative features of the data that require further theoretical explanation.

Our mathematical analysis relied on several simplifying assumptions that could be relaxed in future work. One important simplification was the assumption that constitutively beneficial mutations confer the same characteristic fitness benefit *s*_*b*_. While this caricature is unlikely to be realized in nature, previous work has shown that more general distributions of fitness effects can often be mapped to an equivalent single-*s* model with appropriately chosen values of *s*_*b*_ and *U*_*b*_ (52–54, 56). We therefore expect that most of our conclusions will be robust to relaxations of this particular assumption. Simulations with a more realistic exponential distribution of fitness effects support this intuition (Fig. S3), exhibiting qualitatively similar priority effects as Fig. 2. Another critical assumption was the absence of epistasis, which could influence our current results in a number of important ways. Previous work has shown that global epistasis can cause the fitness effects of new mutations to systematically decline over time as the population fitness increases (74–82). Our analysis shows that this effect would tend to stabilize an existing pair of ecotypes by increasing *T*_sw_ and slowing the rate of cross-invasion — an effect that was previously observed in simulations in Ref. (58) (SI Section 3.5). On the other hand, it is possible that strategy mutations could also alter the supply of new mutations by rewiring an organism’s metabolism, creating new opportunities for compensatory mutation (81). In this case, the different rates of adaptation of the ecotypes could overwhelm the dynamical priority effects we identified above (56). Accounting for this interplay between diversification and evolvability — as well as additional effects like recombination (83–85) — would be a valuable direction for future work.

Finally, many natural microbial communities are considerably more diverse than the two-resource environments we have focused on here. While most of our results continue to hold for larger numbers of supplied resources (SI Section 3), accounting for larger numbers of coexisting ecotypes remains a pressing open challenge (16, 34, 35). We expect that the analytical tools developed in this work — particularly the dynamical establishment picture in SI Section 3.3 — may be useful for extending our current results to the many-ecotype limit as well. However, in this case, the genealogical relationships between ecotypes can be considerably more complex than Fig. 5, and the resulting network of diversification can play an important role in shaping the long-term structure of the ecosystem (34). Future efforts to account for these effects will be critical for understanding how large microbial communities evolve.

## Acknowledgments

We thank Oskar Hallatschek and Daniel Fisher for useful discussions, and J. Lopez, S. Walton, and J. McEnany for comments and feedback on the manuscript. This work was supported in part by the Alfred P. Sloan Foundation (FG-2021-15708), NIH NIGMS Grant No. R35GM146949, and a Terman Fellowship from Stanford University (B.H.G.). B.H.G. is a Chan Zuckerberg Biohub – San Francisco Investigator.

## Author contributions

Conceptualization: D.P.W. and B.H.G.; theory and methods development: D.P.W. and B.H.G..; analysis: D.P.W. and B.H.G.; writing: D.P.W. and B.H.G.

## Code availability

Scripts to generate all simulations and plots are avaiable at https://github.com/DanielWongPGH/consumer_resource_clonal_interference.

## Supplementary Information

**Fig S1.**
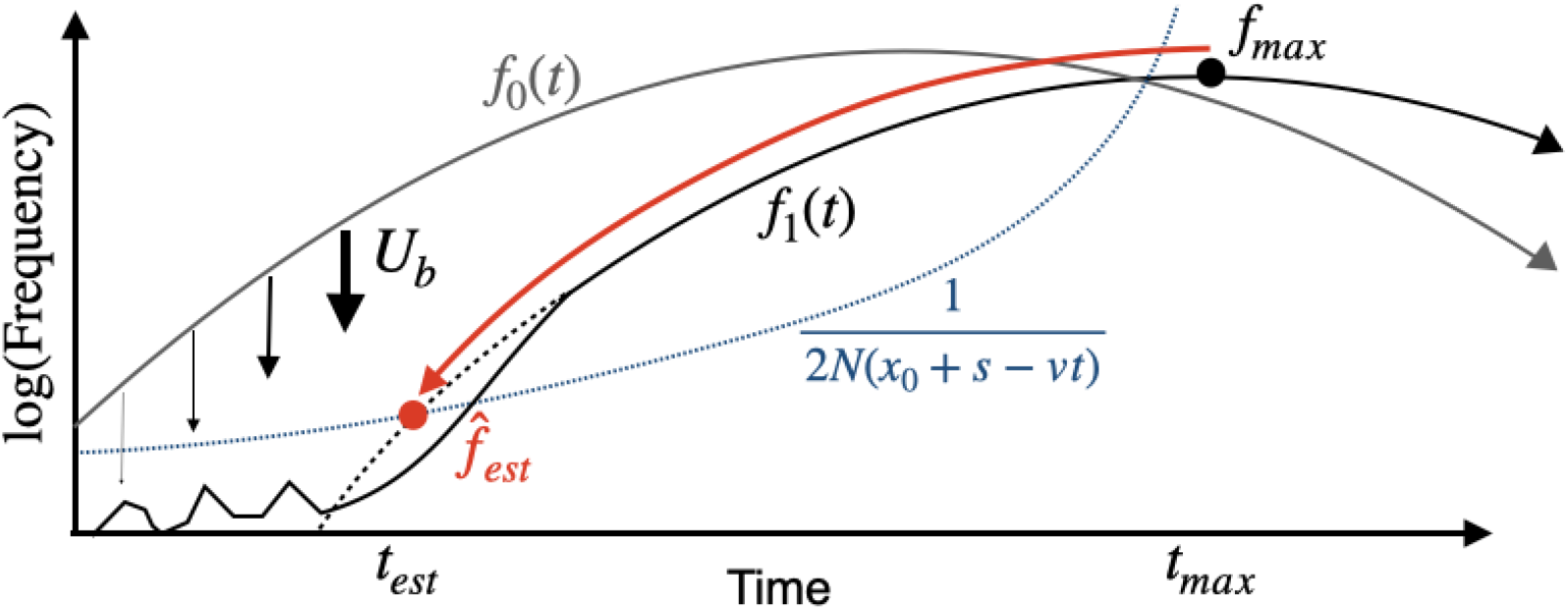
Schematic of the definition of establishment fitness. A lineage *f*_0_ (*t*) with a declining relative fitness *x*_0_ − *vt* produces beneficial mutations that found a new fitness class *f*_1_ (*t*). The establishment fitness *x*_1_ of the new fitness class can then be defined by the relative fitness it would have had at its back-extrapolated establishment time; see SI Sections 3.3 and 4 for more details.

**Fig S2.**
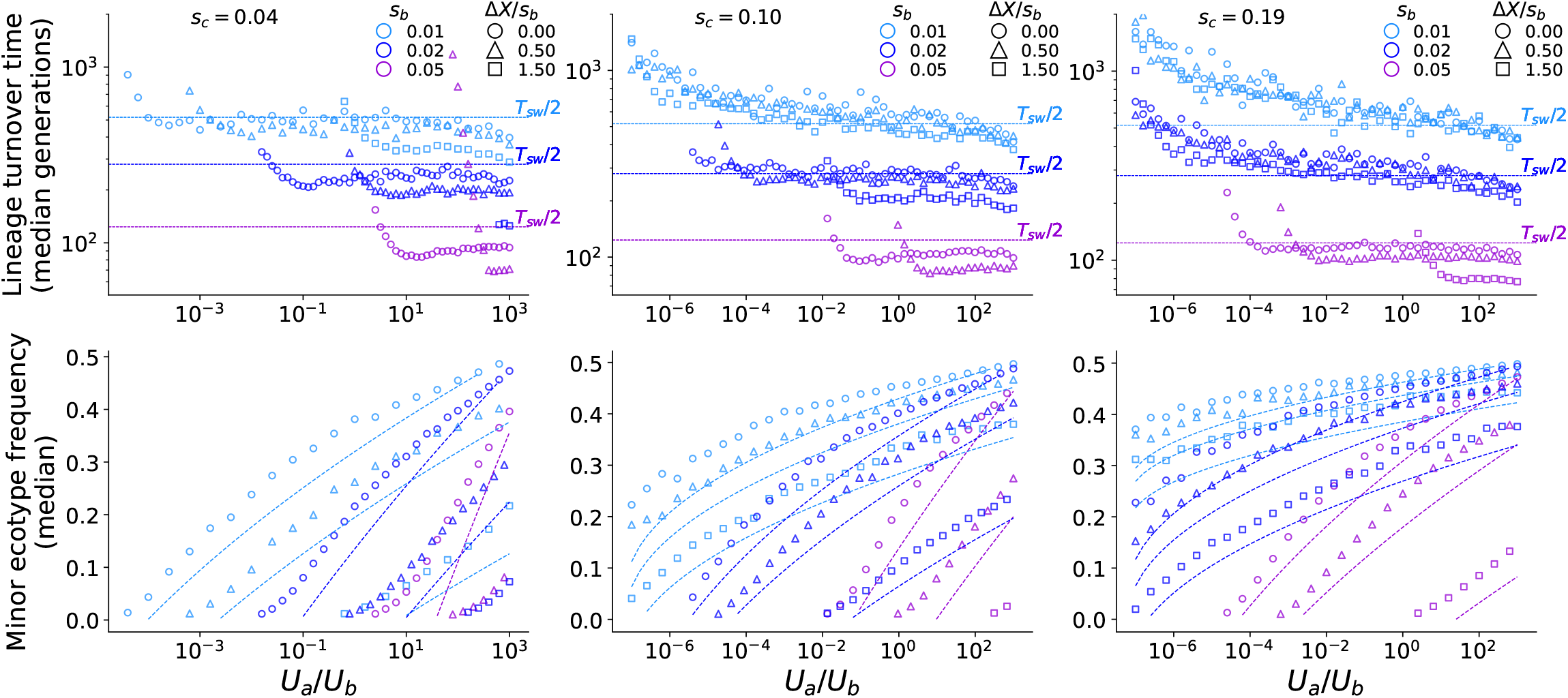
Analog of Fig. 4C,D for strategy mutations with non-zero direct cost *ΔX* < 0. Populations in which strategy mutations incurred variable direct costs of relative strength *ΔX*/*s*_*b*_ were simulated. Ecosystems with differing strengths of ecological feedback B2 are arrayed from left to right. All other parameters as in Fig. 4C,D. At sufficiently strong ecological feedback strength relative to the strength of within-ecotype evolutionary dynamics (by increasing *s*_*c*_ left to right, or decreasing *s*_*b*_ from purple to light blue), direct costs have negligible effect on the typical cross-invasion timescale.

**Fig S3.**
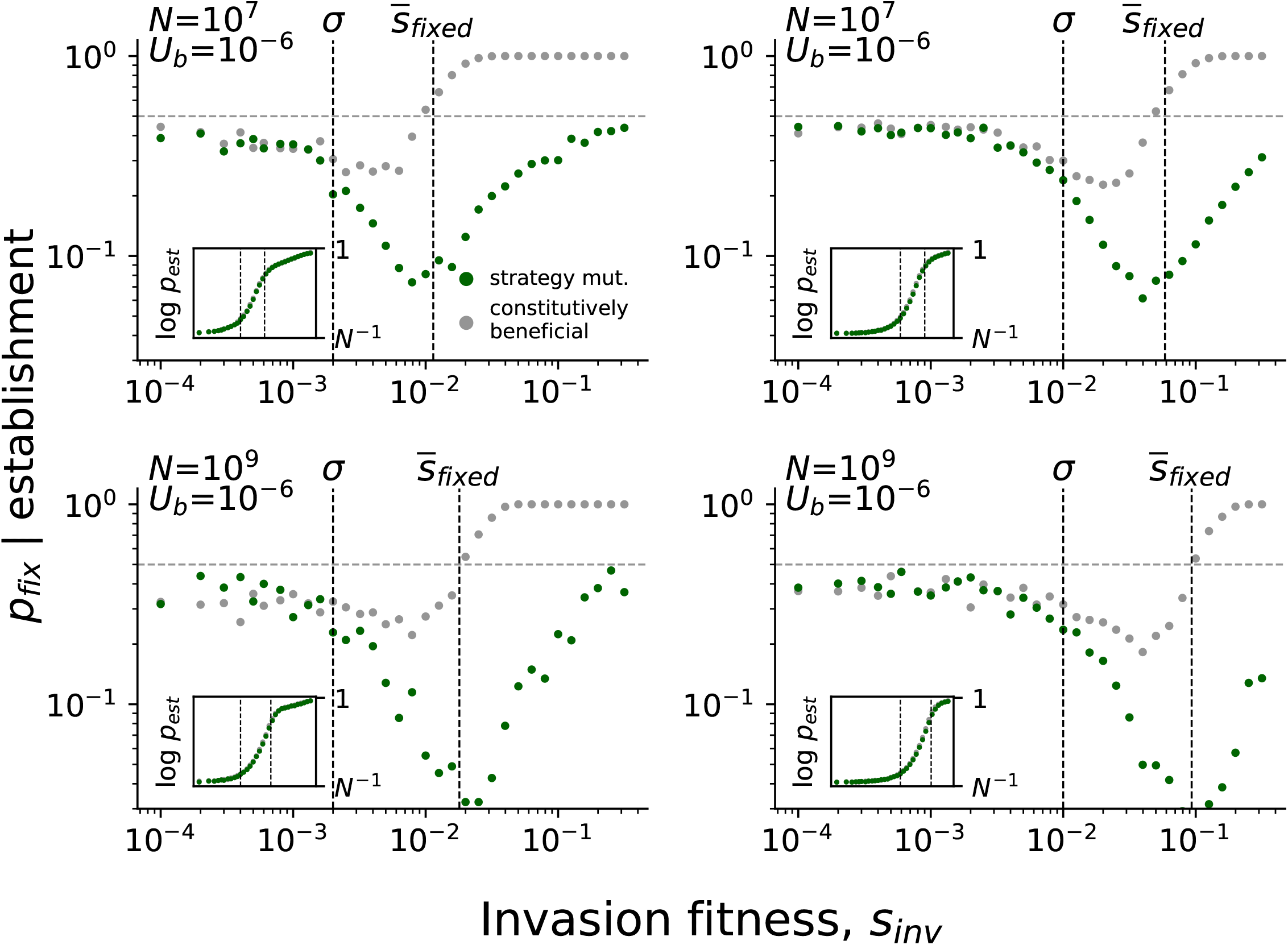
Invasion of strategy mutations in populations with an exponential distribution of general fitnesses. Probability of fixation of a strategy mutation with Δ*X* = 0, *f* ^*^ = 1/ 2, conditioned on establishing (reaching 10% frequency), in clonally interfering populations with an exponential distribution of general fitness effects (DFE) of scale *σ*. Each panel represents a different set of evolutionary parameters (*N*, *U*_*b*_, *σ*), with the mean fixed effect 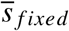 also indicated. To compare to pure strategy mutations (green), conditional probabilities for fitness mutations of equivalent effect Δ*X* = *s*_inv_ are also shown (grey). Insets shows the corresponding establishment probability for strategy and equivalent fitness mutations.

## 1 Model and notation

In this section, we derive the ecological model used in Eq. (1). Our derivation closely follows the ones provided in Refs. (34, 35), which used the same underlying model.

### Consumer-resource model

We consider a population of asexually producing strains *μ*, each of population size *n*_*μ*_. These strains consume ℛ substitutable resources at concentrations 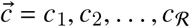 in a chemostat of volume *V*. The number of individuals *n*_*μ*_ belonging to strain *μ* obey the dynamics:

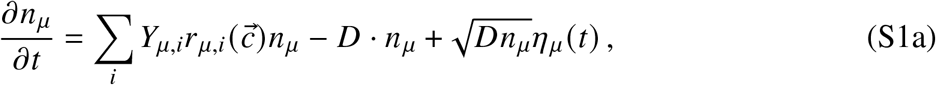

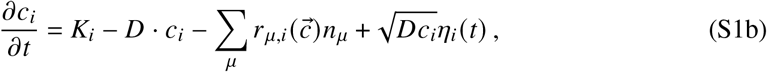

where *D* is the dilution rate, *K*_*i*_ is the input flux of each resource, 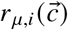 is the total uptake rate of resource *i* by strain *μ, Y*_*μ,i*_ is the yield of resource *i* when converted into biomass by strain *μ*, and *η*_*μ*_ (*t*) and *η* _*j*_ (*t*) are Brownian noise terms (66) with mean zero and variance one. Within this framework, we consider a simplified model where the yields are independent across strains,

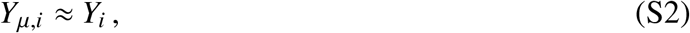

and the uptake function can be decomposed as

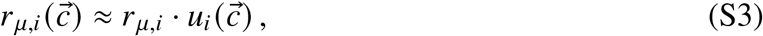

where *r*_*μ,i*_ is a strain-dependent “expression level” and 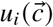 is a corresponding uptake function that is shared across the different strains. We expect that both assumptions will approximately hold in the scenarios we consider in this work, wherein the community is composed of closely-related strains that have recently diversified from a common ancestor.

### Coarse-grained ecological dynamics

Following previous work (34, 35, 58, 59), we consider a limit of Eq. (S1) where the net uptake of each resource is sufficiently fast that the input and uptake terms dominate the resource dynamics in Eq. (S1b). Substituting Eq. (S3) into Eq. (S1), this yields the adiabatic approximation,

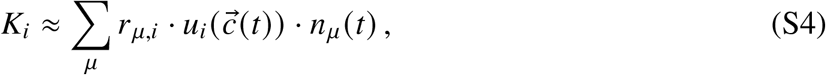

which applies on timescales larger than

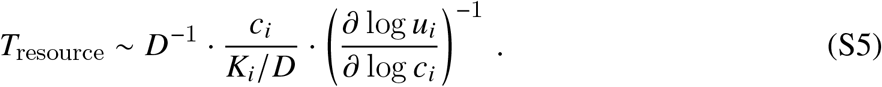

This relaxation timescale will be much shorter than the dilution timescale when *c*_*i*_ is sufficiently small compared to *K*_*i*_ /*D*, which is equal to the steady-state resource concentration that would be attained in the absence of consumption. Ref. (35) showed that this condition will generally hold for small enough values of the dilution rate. Substituting into Eq. (S4) into Eq. (S1a) yields a set of coarse-grained dynamics for the strain abundances alone,

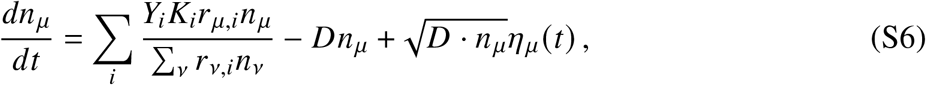

whose the total population size approaches a steady-state value of *N* ≈ ∑_*i*_ *Y*_*i*_*K*_*i*_ /*D* on a timescale of *D*^−1^. Rescaling time in units of *D*^−1^ and defining the scaled variables

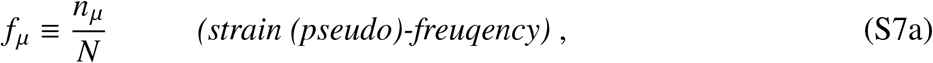

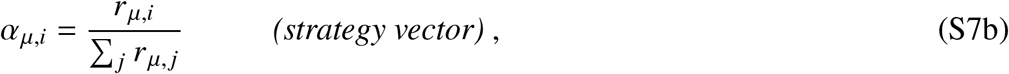

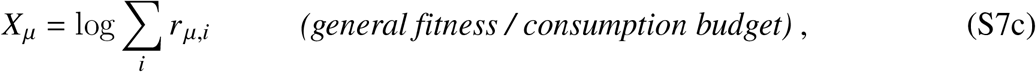

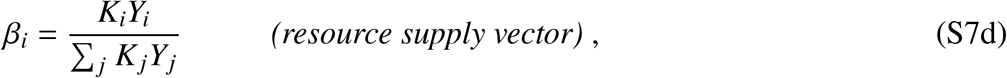

we arrive at the coarse-grained model from Refs. (34, 35, 58, 59):

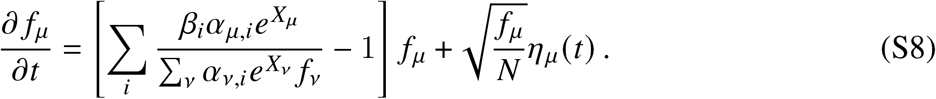

### Limiting form

For most of this work, we will focus on a particular limit of Eq. (S8), where the ecological dynamics take place on timescales much longer than a single generation, and the details of the underlying birth-and-death-process start to become less important. This limiting form is analogous to the diffusion limit of population genetics, and occurs when (i) the maximum growth rates of the different strains (*X*_*μ*_) are sufficiently close to each other (but not identicial), and (ii) the community-averaged-uptake rates 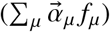 are sufficiently close to the external resource supply vector 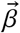. By rewriting both of these quantities in the perturbative form,

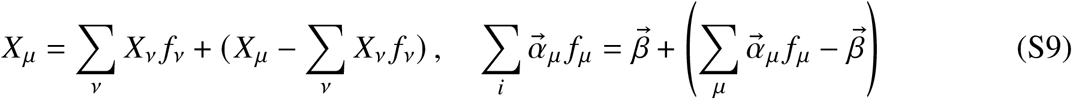

we can expand Eq. (S8) in the limit that |*X*_*μ*_ −∑_*ν*_ *X*_*ν*_ *f*_*ν*_ | ≲ 1 and |*α*_*μ,i*_ *f*_*μ*_ − *β*_*i*_ | ≲*β*_*i*_ to obtain Eq. (1) in the main text,

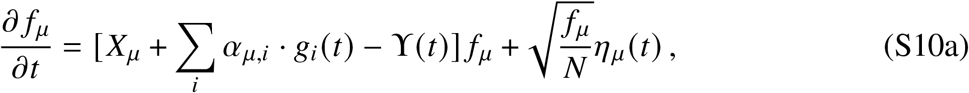

where

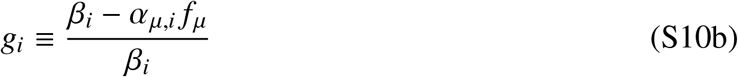

can be interpreted as the local availability of resource *i* and

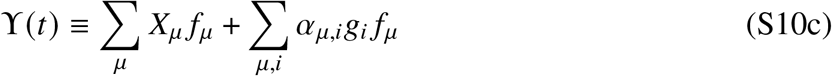

is a gauge term that ensures that the relative frequencies remain normalized. These Lotka-Volterra-like dynamics are equivalent to the “near-ESS limit” of Ref. (34). While Eq. (S10) captures much of the same qualitative as the original model in Eq. (S8), the limiting version in Eq. (S10) tends to admit much simpler analytical expressions, so we will focus on this model for most of our mathematical derivations.

### Incorporating evolution

As described in the main text, we incorporate evolution into the ecological model in Eq. (S10) by allowing the strains to mutate in different ways. Formally, this can be done by adding a mutation kernel to Eq. (S10),

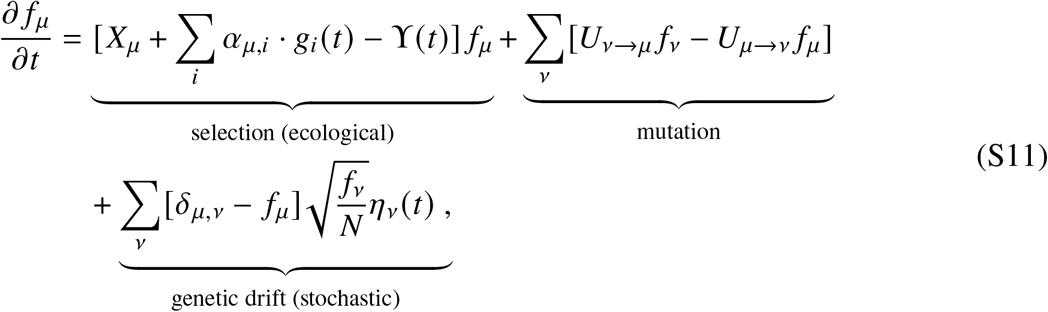

where *U*_*μ* →*ν*_ is the rate of mutation from strain *μ* to strain *ν*. While the ecological dynamics in Eqs. (S8) and (S10) possess a convex Lyapunov function, this is no longer true when mutations are included in Eq. (S11), except in the special case where the mutation kernel is independent of the genetic background (*U*_*μ* →*ν*_ ≈*U*_*ν*_). The resulting dynamics will sensitively depend on the choice of the mutation kernel, which we analyze in SI Sections 2 and 3 below.

## 2 Eco-evolutionary dynamics in the weak mutation regime

Previous studies have analyzed Eq. (S11) in the weak-mutation limit (*NU*_*μ* → *ν*_ →0), where mutations are sufficiently rare that the population approaches an ecological equilibrium long before the next mutation occurs (34, 35). Here we briefly review these results, focusing on the ℛ= 2 case; extensions to larger numbers of resources can be found in Refs. (34, 35).

### Invasion fitness

When a mutation first arises in a resident strain and produces a mutant lineage with phenotype *X* + Δ*X* and 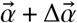, its initial relative growth rate is given by

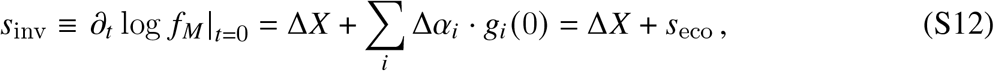

where the value of 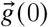 is set by the previous ecological equilibrium. When the resident population contains a single strain, this initial state is particularly simple:

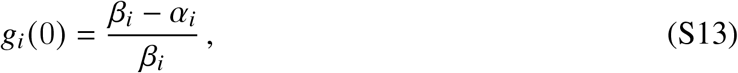

and the invasion fitness is given by

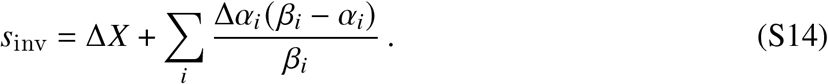

In more general contexts, 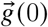 will depend on the community-averaged resource uptake rates in the previous ecological equilibrium, which we discuss in more detail below.

When the mutation first arises, its initial dynamics can be described by the linearized model,

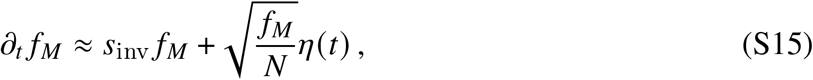

which is valid when the mutant frequency is sufficiently small (*f*_*M*_ *≪*1). These dynamics can be solved by standard methods (86, 87), whose results we briefly summarize here.

Most new mutations will drift to extinction before they reach an appreciable frequency. and the ecological equilibrium of the resident strains will therefore remain unperturbed. However, with probability 2 |*s*_inv_|, the mutant lineage will drift to a size ∼1/ 2*N* |*s*_inv_|, where selection starts to dominate over drift. If *s*_inv_ *<* 0, selection will prevent the lineage from growing much larger than 1/2*N* |*s*_inv_| before it ultimately drifts to extinction. Conversely, if *s*_inv_ *>* 0, then the mutant will start to deterministically expand in the population, and will eventually impact the resource availabilities 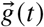. However, the early stochastic dynamics will still be captured by the linearized model in Eq. (S15) whenever the population size is sufficiently large (*N* |*s*_inv_| ≫ 1).

### Ecological equilibrium

If the mutant survives genetic drift, it will start to expand exponentially at rate *s*_inv_ and eventually lead to a new ecological equilibrium. When the resident population is composed of a single strain, these transient dynamics take on a particularly simple form,

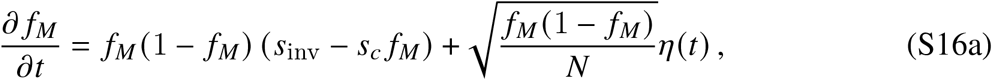

where *s*_*c*_ is defined by

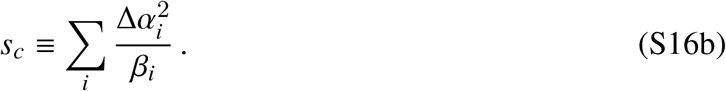

These dynamics reduce to Eq. (S15) when *f*_*M*_ ≪1, while capturing the additional environmental feedbacks that occur when the mutant lineage reaches higher frequencies. The form of this expression shows that the environmental feedbacks will tend to reduce the relative growth rate of the mutant. Whether this reduction is large enough to prevent the mutant from fixing will depend on the relative values of *s*_inv_ and *s*_*c*_. If *s*_inv_ *> s*_*c*_, then the mutant will fix before its relative growth rate is fully reduced to zero. However, if *s*_inv_ *< s*_*c*_, the mutant strain will eventually saturate at an intermediate frequency,

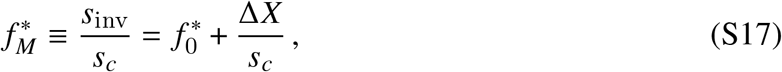

where 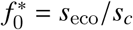. When ℛ= 2, this reduces to the expression quoted in Eq. (3) in the main text. The transient dynamics become more complicated when the initial community is composed of multiple strains. However, in the case of ℛ= 2 resources, at most two strains can coexist with each other at long-times (34, 59, 88). This implies that the equilibrium frequencies are still captured by Eq. (S17) as long as Δ*X* and Δ*α* are defined relative to the phenotype of the coexisting partner. The identity of this surviving strain is uniquely determined by the condition that it has a positive invasion fitness in a population that is entirely composed of the mutant. Substituting Eq. (S17) into Eq. (S10b), one can show that the excess resource availabilities at this multi-strain equilibrium are given by

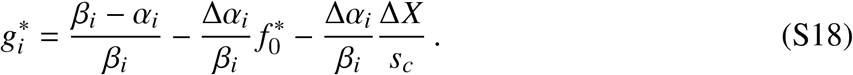

This sets the invasion fitness landscape that applies when the next mutation arises. In the special case of ℛ = 2 resources, this reduces to

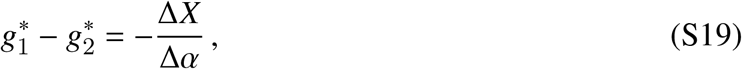

where Δ*α* denotes the resource strategy shift on the first resource.

The transient dynamics in Eq. (S16a) also shed light on the timescales required for equilibration. The deterministic portion of Eq. (S16a) has an implicit analytical solution,

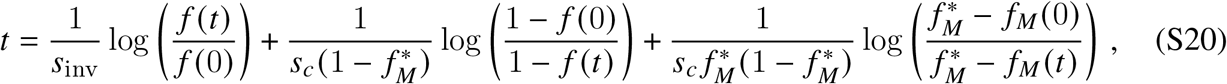

which depends on *s*_inv_, *s*_*c*_, and 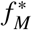. This expression shows that it takes 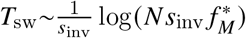 generations for the mutant to grow from an initial frequency of ∼ 1/*Ns*_inv_ to an appreciable fraction of 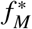, and then another ∼1 /*s*_*c*_ generations to stabilize at the equilibrium frequency. Since the time between successful mutations is inversely proportional to *NUs*_inv_, this shows that the weak mutation limit will start to break down whenever *NU* log(*Ns*_inv_) ≳1.

## 3 Eco-evolutionary dynamics in the clonal interference regime

In large populations where mutations occur more frequently (*NU ≫*1), the simulations in Fig. 1 suggest that the population often self-organizes into a smaller number of coarse-grained ecotypes, which are composed of strains that share the same resource uptake strategies (*α*_*μ*_) but have different genotypes (*μ*) and fitnesses (*X*_*μ*_). To analyze these dynamics, it is useful to define a corresponding set of ecotype frequencies,

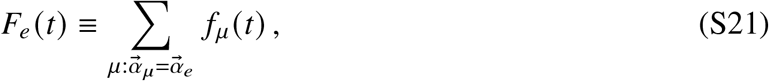

where 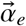 denotes the resource uptake strategy of ecotype *e*. By summing over the strain frequencies in Eq. (S11), one finds that the ecotype frequencies obey a related equation,

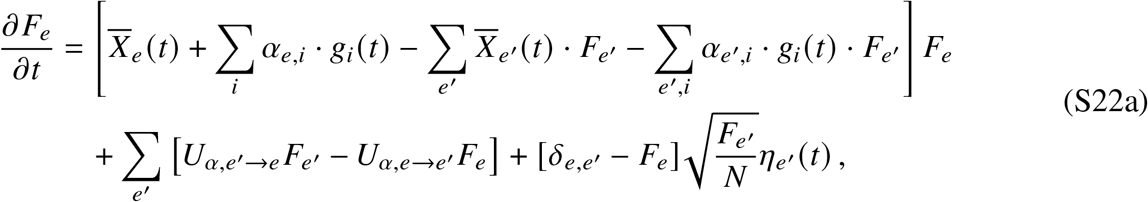

Where 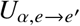 is the rate of strategy mutations from ecotype *e* to *e*,,

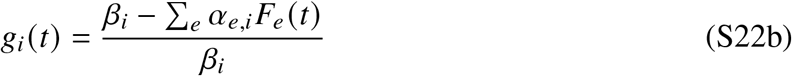

is the excess resource availability, and

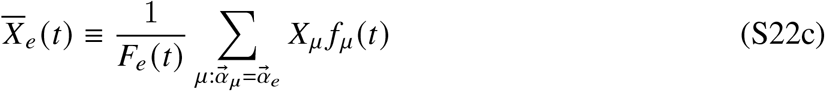

is the average *X* coordinate (e.g. metabolic budget) of ecotype *e*. In the simplest case of two ecotypes, this reduces to Eq. (6) in the main text. Eqs. (6) and (S22) show that the ecotype frequencies follow a structurally similar set of dynamics as the underlying strain frequencies in Eq. (S11), except that the fixed parameters *X*_*μ*_ are now replaced with the time-dependent variables 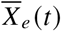. However, since these time-dependent quantities depend on the internal composition of each ecotype, the dynamics in Eq. (S22) do not fully close, and will not generally possess an analogous ecological equilibrium when constitutively beneficial mutations are common (*NU*_*b*_ *≳* 1).

In the following sections we develop a theory for the intra-ecotype dynamics that set 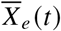, and how they interact with the inter-ecotype dynamics in Eq. (S22). These sections follow the organization of the main text, starting from the dynamics of a single ecotype and progressively introducing more complexity.

### 3.1 Evolutionary dynamics of a single ecotype

When the population is composed of a single ecotype, the resource availabilities in Eq. (S11) cancel out with each other, and the dynamics reduce to

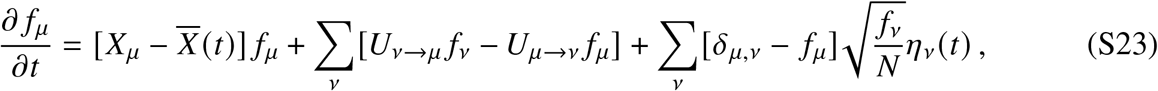

where 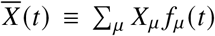 is the mean fitness of the population. This equation has the same form as earlier “non-ecological” models of clonal interference that have been studied in previous work (50–55). We briefly summarize these results for the single-*s* model in Fig. 1, focusing on the empirically relevant regime where *s*_*b*_ *≫U*_*b*_ *≫*1 *N*. We will then extend these results to ecological contexts in the following sections.

When beneficial mutations are common (*NU*_*b*_ ≫1), previous work has shown that the resident ecotype can be described as a traveling “fitness wave”, with a characteristic shape *ρ x* that moves toward higher fitness at a steady-state rate 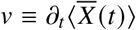. In our parameter regime of interest, this relative fitness distribution is well-approximated by a truncated Gaussian shape,

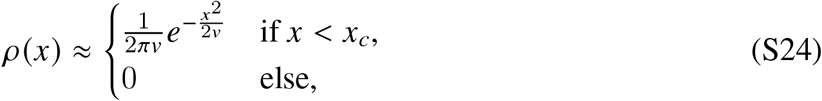

where 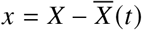 denotes the relative fitness, and *x*_*c*_ denotes the location of the fittest individuals that are currently alive in the population (53–55). These highly fit individuals in the nose of the distribution will eventually give rise to further mutations, which ultimately allows the distribution in Eq. (S24) to steadily move to higher fitness values.

The steady-state values of *v* and *x*_*c*_ can be determined by tracking the long-term fate of a lineage founded by an individual with relative fitness *x*. The descendants of this lineage can be divided into “fitness classes” *f*_*k*_, where *k* ≥ 0 denotes the number additional beneficial mutations that they have acquired. In order to fix, the focal lineage must repeatedly seed additional fitness classes, and outrun the steady advance of the bulk population. Previous work (53–56) has shown that this fixation probability can be calculated from a branching process approximation to Eq. (S23), where the mean fitness is dominated by the background population 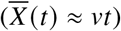. When *x > x*_*c*_ −*s*_*b*_, this fixation probability is well-approximated by the piecewise form,

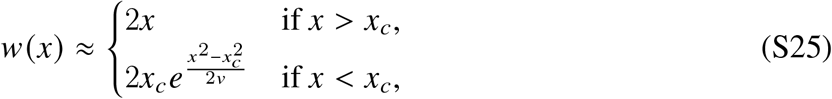

with *x*_*c*_ given by

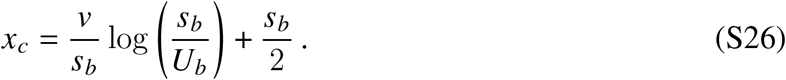

We can use this result to calculate the fixation probability of a new mutation (*X* → *X* +*s*), by averaging over the genetic backgrounds it could have arisen on:

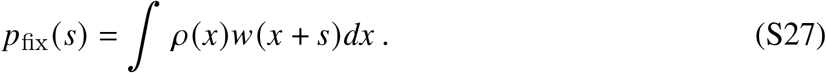

The requirement that *v* = *NU*_*b*_ · *p*_fix_(*s*_*b*_) · *s*_*b*_ yields a second self-consistency relation,

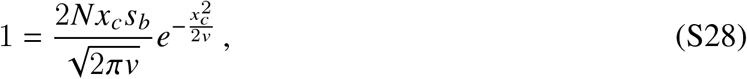

which can be combined with Eq. (S26) to solve for the rate of adaptation in terms of *N, s*_*b*_, and *U*_*b*_.

The leading order contributions are given by

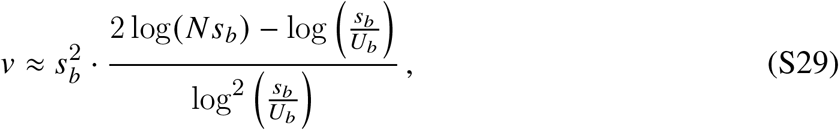

where the weak dependence on *N* and *U*_*b*_ reflects the fact that most new mutations are lost to clonal interference.

#### Conditions of validity

The results in Eqs. (S24)-(S29) are valid in the asymptotic regime where 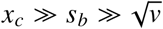. The first condition (*x*_*c*_ ≫*s*_*b*_) implies that the fittest individuals are substantially more fit than the majority of the population – i.e. that clonal interference is common. The second condition 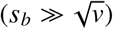 implies that the majority of the population is concentrated within one mutation of the mean fitness. Using the approximate expressions for *x*_*c*_ and *v* in Eqs. (S26) and (S29), we can obtain an analogous set of conditions on the underlying parameters *N, s*_*b*_, and *U*_*b*_:

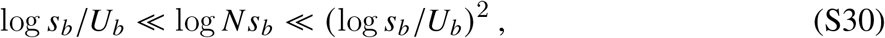

These conditions are approximately satisfied for many experimental microbial populations, where log_10_ *s*_*b*_ ≃ −2 ± 1, log_10_ *U*_*b*_ ≃ −6 ± 1, log_10_ *N* 7 ± 1 (52). However, since the relevant quantities are all logarithmic functions of *N, s*_*b*_, and *U*_*b*_, convergence to this limit can be slow in practice. Nevertheless, we generally find that the analytical results provide a good match to simulations even when *x*_*c*_ /*s*_*b*_ and 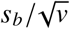 are only modestly large (e.g. ≥ 2). We therefore continue to assume that *x*_*c*_ ≫ *s*_*b*_ and 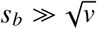 in all of our subsequent calculations, since it allows us to simplify many of our key results.

#### Fluctuations in the speed of the wave

The rate of adaptation *v* is only an average, for two reasons. First, on short timescales, the discreteness of the underlying fitness classes causes the mean fitness to increase in a step-like manner. Previous work has shown that when 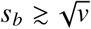, the dominant fitness class transitions from *k* to *k* +1 on a timescale of ∼1 *s*_*b*_ generations, followed by a longer period of stasis before the next transition occurs roughly ∼*s*_*b*_ /*v* generations later (52). In addition to this effective periodicity, the inherent randomness in the timing of new mutations in the nose of the fitness distribution generates corresponding fluctuations in the speed of the wave that manifest over longer timescales (52, 54). We consider these fluctuations in more detail in SI Section 3.5, where they play an important role in driving the long-term competition between two established ecotypes. However, in most other contexts, we find that the deviations from a constant rate of adaptation have a negligible impact on our main results, so we will often invoke this assumption for simplicity.

### 3.2 Establishment of a new ecotype

Building on the results above, we next consider the invasion of a new ecotype (*α*_*M*_) created by a resource strategy mutation in a population with a single resident ecotype 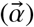. In this case, the resource availabilities in Eq. (S22b) reduce to

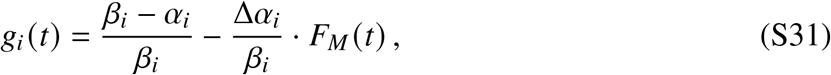

where 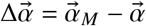 and *F*_*M*_ (*t*) is the total frequency of the mutant ecotype. We assume that when the strategy mutation arises, the population has already been evolving for a sufficient amount of time that the fitness distribution of the resident ecotype has converged to the traveling wave form in Eq. (S24). The new ecotype will therefore arise on a genetic background with relative fitness drawn from *ρ* (*x*), and its descendants will compete with the resident ecotype as they both acquire further mutations. In the absence of additional strategy mutations, the descendants of the initial strategy so that the total frequency of the invading ecotype can be written as *F*_*M*_ (*t*) = ∑ _*k*_ *f*_*M,k*_ (*t*), then the mutation can be fully characterized by the number of additional fitness mutations that they have acquired. If we let *f*_*M,k*_ (*t*) denote the total frequency of mutant individuals with *k* such mutations, so that the total frequency of the invading ecotype can be written as *F*_*M*_(*t*)= ∑_*k*_ *J*_*M,k*_ (*t*), then th dynamics of the invading ecotype can be formally described by a mesoscopic version of Eq. (S22),

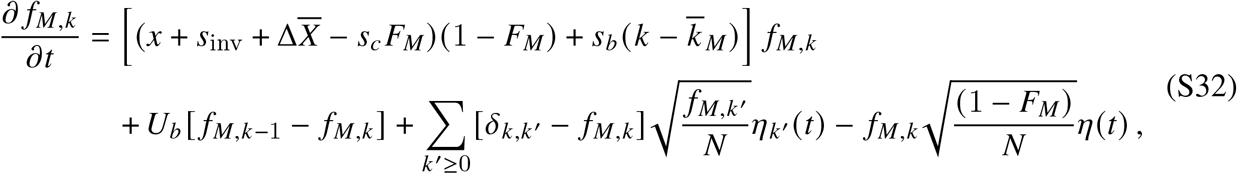

where 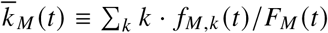 is the average value of *k* in the invading ecotype, 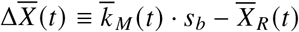 is the difference between the average *X* coordinates of the resident and invading ecotypes, and *s*_inv_ and *s*_*c*_ are the same as in the weak-mutation limit in Eq. (S16). For concreteness, we assume that the 0 *< s*_inv_ *< s*_*c*_, so that the strategy mutation would normally coexist with its parent in the absence of clonal interference. We can then ask how the additional terms in Eq. (S32) alter the dynamics of diversification in Eq. (S16).

The key approximation we will make is that during the initial establishment of a new ecotype, the resident population will continue to be well-described by the traveling wave theory in Eq. (S24). This allows us to replace the time-varying quantity 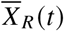 with the deterministic function 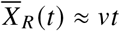, yielding a closed model,

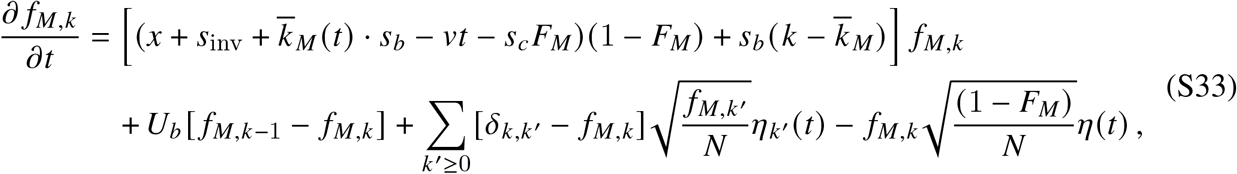

which only depends on the mutant fitness classes *f*_*M,k*_ (*t*). This deterministic approximation will hold until the established ecotype grows to a large enough size that it influences the dynamics of 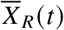; we treat these interactions more explicitly in SI Section 3.5 below.

The simplified model in Eq. (S33) is still quite complicated, since it includes both stochastic and nonlinear terms. Borrowing intuition from the non-ecological case, we expect that the intermediate fate of the invading ecotype will be primarily determined when it is still at a low frequency in the population (*F*_*M*_ 1). In this limit, Eq. (S33) reduces to the linearized model,

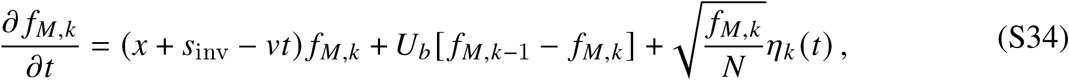

which is equivalent to a non-ecological mutation with an initial relative fitness of *x* + *s*_inv_. This suggests that the initial establishment probability of an invading ecotype — i.e. the probability that it initially outraces the resident and reaches a macroscopic frequency in the population — can be approximated by Eq. (S27) above, but with *s* = *s*_inv_. This yields Eq. (5) in the main text, which we reproduce here for convenience:

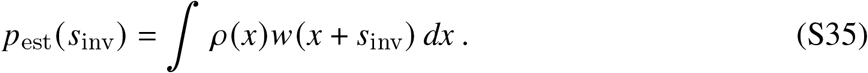

This approximation is borne out in simulations (Fig. 2B), though we note that it somewhat underestimates our operational definition of establishment in Fig. 2 (i.e. reaching *F*_est_ ≈ 10% frequency) when 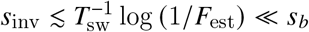. This discrepancy is an artifact of our operational definition of establishment: it reflects the growing contributions of nearly neutral trajectories that never truly establish. Since these contributions become negligible for larger fitness benefits 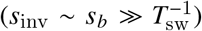, we will omit them for simplicity, and use Eq. (S35) as our new operational definition of establishment.

While the initial dynamics of the invading ecotype can be summarized by Eq. (S35), its longer-term trajectory will sensitively depend on the behavior of 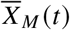 at later times (Eq. S22). This requires additional information about the establishment process in Eqs. (S33) and (S34) that is not directly encoded in Eq. (S35). In the following sections, we present an alternative framework for predicting the establishment probability of a new mutation that provides this additional dynamical information. This framework is conceptually similar to the genetic draft models in Refs. (89, 90), but recast in (continuous) relative fitness space. It is also closely related to the front fluctuation analysis in Ref. (54), and leverages much of the same mathematical machinery. While formally equivalent to the establishment probability in Eq. (S35), this dynamical picture of the establishment process will be particularly useful for modeling the trajectories of invading ecotypes in SI Section 3.4.

### 3.3 Dynamical establishment picture

To streamline notation, we will suppress the *M* index in Eqs. (S32-S34). We will then focus on the dynamics of successive fitness classes *f*_*k*_ (*t*), which are populated in a sequential manner. The behavior of the *k* = 0 class is particularly simple, since it was founded by a single mutation at time *t* = 0. While the frequency of the invading ecotype remains rare (*F*(*t*) ≪ 1), the dynamics of the founding class in Eq. (S34) reduce to the same form as Eq. (S15), but with the time-varying invasion fitness,

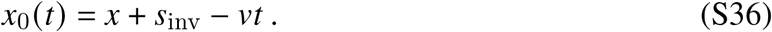

Provided that 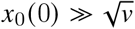, this founding clone will survive genetic drift with probability *p*_est_ ≈ 2*x*_0_(0), and will then begin to grow as

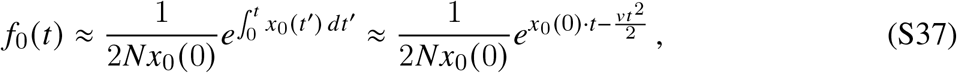

before reaching a maximum size of *f*_max_ ≈ exp *x*_0_ (0) ^2^ / 2*v*) / 2*Nx*_0_ (0) when *t ≈ x*_0_ (0) / *v* (Fig. S1). Comparing this expression with Eq. (S28), we can see that a key transition occurs when *x*_0_ 0 is close to *x*_*c*_. If *x*_0_ 0 is larger than *x*_*c*_, then the founding clone will eventually grow to 𝒪 (1) frequencies and will need to be modeled by the full dynamics in Eq. (S33); we consider this case in more detail in SI Section 3.4 below. On the other hand, if *x*_0_ (0) *< x*_*c*_, then the founding clone will remain rare throughout its entire lifetime, and additional fitness mutations are necessary for the invading ecotype to reach large frequencies.

#### Initial fitnesses of successful founding clones

The relative contributions of these two scenarios can be quantified by revisiting the integral in Eq. (S35). The posterior distribution for initial relative fitness of an established mutation is proportional to the integrand in Eq. (S35),

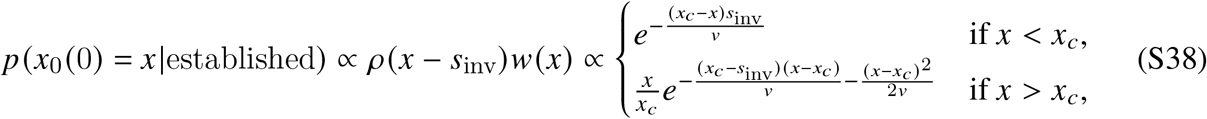

where the second line follows from substituting the analytical expressions in Eqs. (S24) and (S25). The shape of this distribution strongly depends on how *s*_inv_ compares to *x*_*c*_. When *s*_inv_ ≳ *x*_*c*_ */* 2, the distribution of successful *x*_0_ 0 values is dominated by the region where *x*_0_ (0) *> x*_*c*_. For these mutations, the founding strain of the new ecotype will typically be the first to grow to reach large frequencies. On the other hand, when *s*_inv_ ≲ *x*_*c*_ 2, the distribution of of successful *x*_0_ (0) values is dominated by the *x*_0_ (0) *< x*_*c*_ region of Eq. (S38), with a typical value of *x*_*c*_ − *x*_0_ (0) *v s*_inv_. For such moderate effect mutations — which includes all those with fitness benefits comparable to the typical driver mutation *s*_*b*_ — additional fitness classes are typically required for the invading ecotype to reach large frequencies.

#### Establishment of subsequent fitness classes

Returning to Eq. (S34), the low-frequency dynamics of the *k* = 1 class are given by

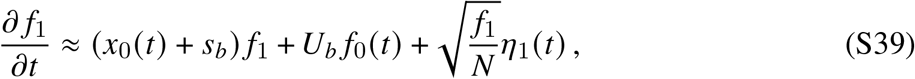

where *f*_0_ (*t*) is given by Eq. (S37). The key difference from the founding *k* = 0 class is that *f*_1_ (*t*) must be seeded by future mutations that originate from *f*_0_ (*t*). The establishment of a growing population fed by mutations from another exponentially growing population has been analyzed in a number of previous studies (52, 89, 91, 92). In SI Section 4, we generalize these earlier results to incorporate the additional *v* dependence in the continuous fitness model in Eq. (S39).

Following Ref. (52), we can write the long-time dynamics of *f*_1_(*t*) in the approximate form

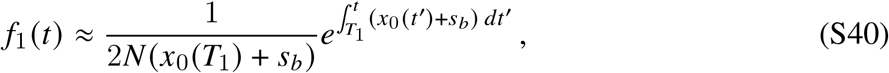

where *T*_1_ is a random variable representing the *establishment time* of the *k* = 1 class (Fig. S1; SI Section 4). Strictly speaking, this approximation is only accurate for *t* ≳ *T*_1_ + 𝒪 (1 / *s*_*b*_), but this will be sufficient to capture the dynamics of *f*_1_ (*t*) on the timescales of interest where it starts producing further mutations. In our continuous fitness model, Eq. (S40) further reduces to

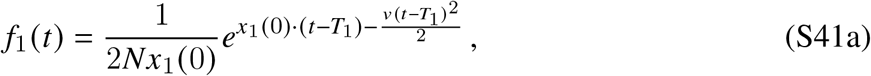

where we have a defined a new random variable,

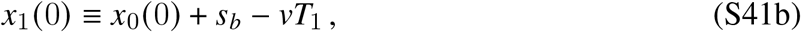

which corresponds to the instantaneous relative growth rate of *f*_1_ (*t*) near its backward-extrapolated establishment time (Fig. S1). We will refer to this quantity as the *establishment fitness* of *f*_1_ (*t*).

We note that the trajectory in Eq. (S41) has the same functional form as the founding mutation in Eq. (S37), but with *x*_0_ (0) → *x*_1_(0), and time shifted forward by an amount *T*_1_. As a result, the long-term dynamics of the *k* = 1 class will be similar to those of the *k* = 0 class above, reaching 𝒪 (1) frequencies if *x*_1_ (0) *> x*_*c*_, and remaining small if *x*_1_ (0) *< x*_*c*_. This analysis can be repeated for subsequent fitness classes with progressively larger values of *k*. Each successive class *f*_*k*_ (*t*) will follow the same self-similar form as Eq. (S41),

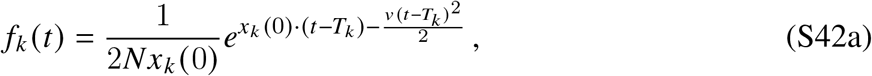

with a corresponding establishment fitness

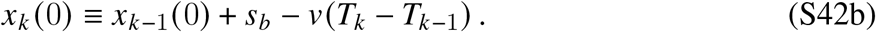

This fitness-class decomposition shows that the establishment of the entire lineage requires the production of at least one fitness class with an establishment fitness larger than *x*_*c*_.

#### Distribution of successive establishment fitnesses

The key remaining question is how to predict *x*_*k*_(0) given the previous values of *x*_*k* −1_ (0) and *T*_*k* − 1_. We can calculate the distribution of *x*_*k*_ (0) from the corresponding distribution of *T*_*k*_, focusing on our parameter regime of interest where |*x*_*k* − 1_(0) *x*_*c*_ − *x*_*c*_ | (SI Section 4). The establishment time distribution has the same skewed form as in the *v* = 0 case in Ref. (52). Most of the probability is concentrated near the average value,

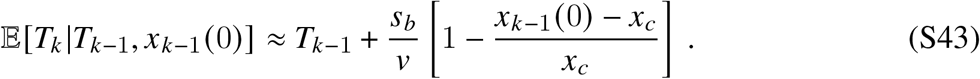

Substituting Eq. (S43) into Eq. (S42b) yields a corresponding recursion for the establishment fitness,

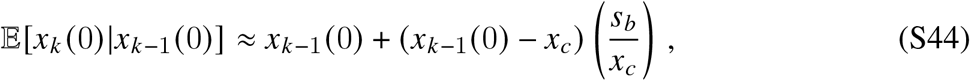

whose solution in terms of *x*_0_(0) is given by

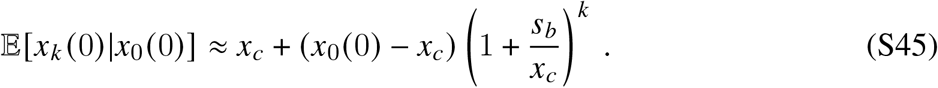

The resulting dynamics exhibit a bifurcation depending on the relative values of *x*_0_ (0) and *x*_*c*_. If *x*_0_(0) *> x*_*c*_, then the expected establishment fitnesses of future fitness classes will tend to grow even larger in each successive iteration; conversely, *x*_*k*_ (0) will tend to shrink in successive classes if *x*_0_ (0) *< x*_*c*_. These selection-like biases are functionally equivalent to the genetic draft effects described in Refs. (89, 90), but now recast in relative fitness space rather than as frequencies within the nose of *ρ* (*x*). Similar results were also obtained for the fluctuations in the wavefront of *ρ* (*x*) in Ref. (54) using a conceptually similar approach. In this case, *x*_*c*_ corresponds to the special point where the establishment fitness of future fitness classes is approximately conserved over time; this is consistent with the intuitive definition of the nose of the steady-state distribution in Eq. (S24).

The full distribution of *x*_*k*_ (0) in SI Section 4 shows that the typical fluctuations around the average values in Eq. (S44) are small provided that |*x*_*k*_ (0) *x*_*c*_ | ≳ *v* /*x*_*c*_. This implies that the average is often a good approximation for the realized sequence of *x*_*k*_ (0) values. However, in each successive fitness class, there is also a small probability that an anomalously early mutation produces a correspondingly earlier establishment time, and a therefore a substantially higher establishment fitness (Δ*x*_*k*_ (0) *> v* / *x*_*c*_), corresponding to a *>e*-fold total frequency change ∼*x*_*c*_ / *v* generations later (52, 54, 89, 91). The analysis in SI Section 4 shows that the probability density of these jackpot events is given by

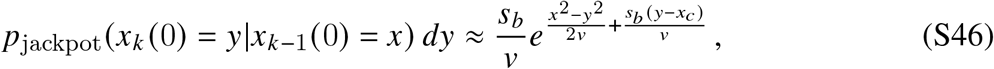

which applies for *v* / *x*_*c*_ ≲ − *x*_*k*_ (0) − *x*_*k* − 1_ (0) ≲ *s*_*b*_. While the total probability of a jackpot is small (*s*_*b*_ *x*_*c*_ 1), they nevertheless play a crucial role in allowing mutations with *x*_0_ 0 *< x*_*c*_ to establish, since this is the only way to reverse the inevitable decline in Eq. (S44) and produce a subsequent fitness class with *x*_*k*_ (0) *> x*_*c*_.

#### Establishment probability of the entire lineage

One can combine the results in Eqs. (S44-S46) to obtain an expression for the establishment probability of the entire lineage, starting from an established founding clone with an initial relative growth rate of *x*_0_ (0) = *x*. If *x > x*_*c*_, this establishment probability is equal to one, since the founding clone (and any subsequent fitness classes) are guaranteed to reach high frequencies. For lower initial fitnesses, the establishment probability can be expressed as a recursion,

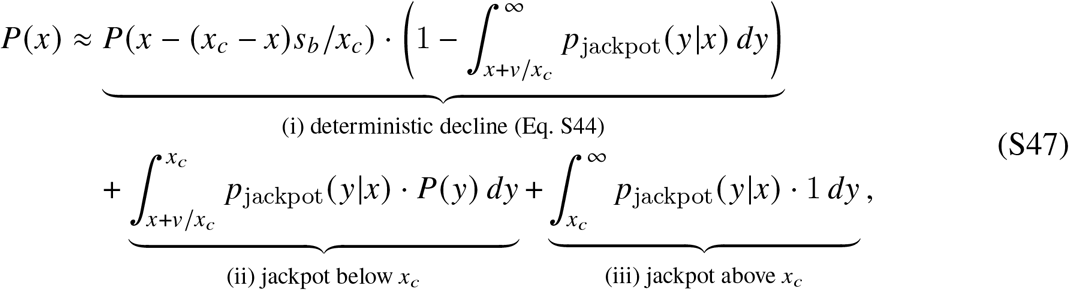

which sums over the different possibilities for the establishment fitness of the next class. This recursion contains three terms corresponding to (i) the steady decline of *x*_*k*_ (0) under the deterministic dynamics in Eq. (S44), as well as rare jackpot events that land both (ii) below and (iii) above *x*_*c*_.

One can check that 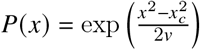 is a solution to Eq. (S47) when *x < x* ; this matches the previous result for *w* (*x*) in Eq. (S25) after accounting for the establishment of the founding clone. In this case, the alternative derivation in Eq. (S47) now demonstrates that (i) the precipitating event for the establishment of the entire lineage is the production of a fitness class with *x*_*k*_ (0) *> x*_*c*_, and (ii) that the posterior distribution of the first such *x*_*k*_ (0) value is given by

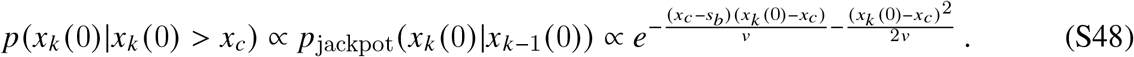

Equation (S48) also approximately matches the posterior distribution of *x* (0) in Eq. (S38) when *s*_inv_ ≲ *x*_*c*_ / 2. We can therefore combine these expressions to obtain the distribution of successful *δ* values listed in Eq. (10) in the main text. We will use this result to predict the frequency trajectory of the established ecotype in SI Section 3.4 below.

Finally, we note that the decomposition in Eq. (S47) also shows that the smaller jackpot events that land below *x*_*c*_ — while not directly leading to establishment on their own — still play a dominant role in setting the overall establishment probability *P*(*x*). By comparing the magnitudes of the two jackpot terms, we see that a successful lineage will typically undergo 𝒪 (*x*_*c*_/*s*_*b*_) intermediate jumps to fitnesses of order *x*_*c*_ − 𝒪 (*v*/*s*_*b*_) before producing a fitness class with *x*_*k*_ (0) *> x*_*c*_.

#### Validity of approximations

In addition to the asymptotic requirements that we have already discussed, namely 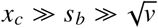 and |*x*_*k*_ 0 − *x*_*c*_| ≪ *x*_*c*_, our results also depend on the validity of the linear fitness class model in Eq. (S34). This model will eventually break down for established lineages, since at least one of the underlying fitness classes will eventually reach a size where they start to impact the surrounding population. However, our dynamical picture shows that these feedbacks have a negligible impact on the underlying probability of establishment, since all fitness classes prior to the one with *x*_*k*_ (0) *> x*_*c*_ are guaranteed to remain at low frequency. This provides a more detailed justification for the assumptions made in previous studies (53–56, 93), which used the branching process model in Eq. (S34) to predict the fixation probabilities of a wide variety of mutations.

### 3.4 Invasion trajectory of a new ecotype

Once an ecotype generates a fitness class with *x*_*k*_ (0)*> x*_*c*_, it will eventually increase in frequency to the point where it will start to impact the surrounding population, and Eq. (S34) will break down. We now consider the invasion trajectories of these initially successful ecotypes, by returning to the full dynamics in Eqs. (S32) and (S33). Without loss of generality, we will assume that the *k* = (0) class is the first one with *x*_*k*_ 0 *> x*_*c*_; the same math will apply in other contexts if we treat the first class with *x*_*k*_(0) *> x*_*c*_ as the effective founder.

We begin by focusing on the initial phase of the invasion trajectory, where the rate of adaptation of the resident ecotype remains close to its initial value 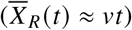, and the fitness class dynamics satisfy Eq. (S33); the dynamics on longer timescales will be treated in SI Section 3.5 below. Summing over the fitness classes in Eq. (S33), the total frequency of the invading ecotype satisfies

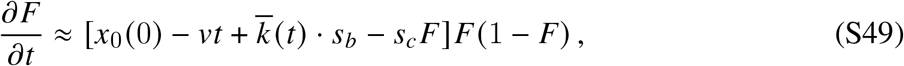

where 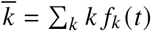 is the average number of additional mutations that have accumulated since the founding class. These dynamics are similar to the weak-mutation version in Eq. (S16a), except that we now have an additional time-dependent term, 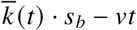, that accounts for the ongoing adaptation of the two ecotypes. We can obtain an approximate analytical solution to Eq. (S49) using asymptotic matching, by dividing the resulting trajectory into distinct phases. For simplicity, we will assume that 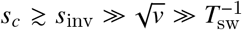, since this includes the portions of Fig. 2 where most of the interesting behavior occurs.

#### Phase 1: Expansion of the founding clone

In the initial phase of each trajectory, the invading ecotype is numerically dominated by its founding class, so that 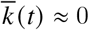, and Eq. (S49) reduces to

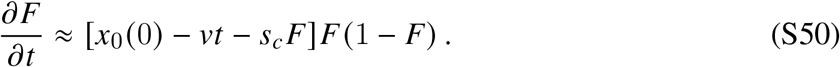

At early times, the higher-order terms in Eq. (S50) can be treated as a small correction, and the ecotype frequency reduces to Eq. (S37):

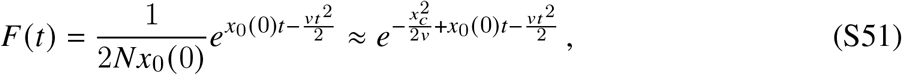

where we have used Eq. (S28) to approximate log *Nx*_*c*_ by 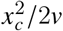.

We can use Eq. (S51) to calculate the total time *T* ^*^ that it takes for *F* (*t*) to reach a size where the ecological feedbacks become important. We denote this critical size by

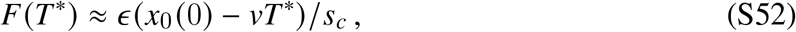

where *ϵ* ≪ 1 is an arbitrary small parameter that is chosen to be larger than any of the other small parameters in our model. Since we have assumed that *s*_inv_ *< s*_*c*_, it will always be the case that *F* (*T* ^*^) ≪ 1, so Eq. (S51) will still apply. Solving for *T* ^*^ then yields

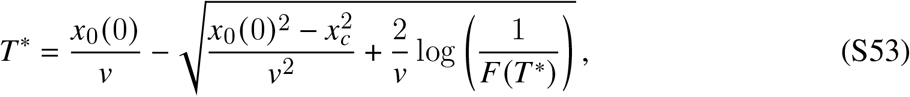

which reduces to the simpler version in Eq. (7) in the main text when 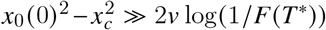.

We note that for |*x*_0_ (0) − *x*_*c*_ | ≪ *x*_*c*_, this critical time *T* ^*^ is smaller than the characteristic sweep time *T*_sw_ ≈ *x*_*c*_ / *v* of the resident ecotype. This implies that the residual growth rate of the founding clone,

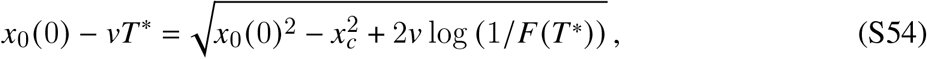

will be larger than its growth advantage relative to the nose of the resident ecotype – often by substantial amounts. This excess relative fitness will drive the overshooting dynamics that we now consider in the next phase of the trajectory.

#### Phase 2: Rapid ecological feedback

At longer times (*t > T* ^*^), the higher-order terms in Eq. (S50) will begin to slow further growth, and Eq. (S51) will break down. The resulting dynamics will depend on the magnitude of the residual growth rate *x*_0_(0)− *vT* ^*^ relative to the ecological restoring force *s*_*c*_.

##### Small invaders

If *x*_0_ (0) − *vT* ^*^ *s*_*c*_, then the ecological feedbacks will become important while the invading ecotype is still at low frequencies (*F* (*t*) ≪ 1). In this case, Eq. (S50) reduces to the simpler form,

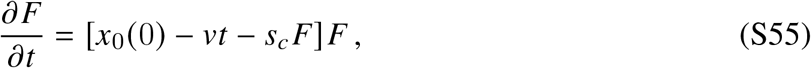

whose solution is given by

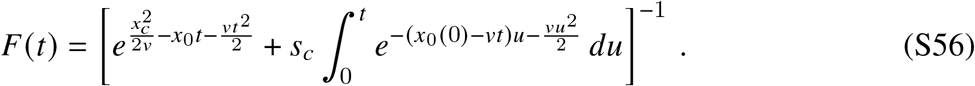

This expression can be interpreted as the harmonic mean of the unconstrained trajectory in Eq. (S51) and a new local equilibrium,

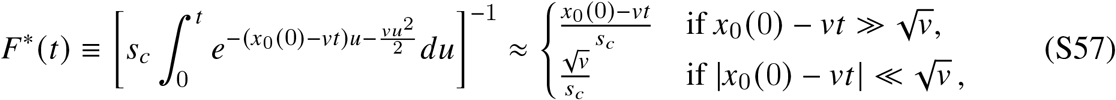

which is the time-dependent analogue of *f* ^*^ in SI Section 2. Eq. (S56) shows that the invading ecotype will exhibit a peak soon after *t* ≈ *T* ^*^, before saturating at the declining local equilibrium *F*^*^(*t*) at longer times.

##### Large invaders

For larger values of *s*_inv_, the residual growth rate *x*_0_ (0) − *vT* ^*^ will remain of order *s*_*c*_, and it will not always be possible to drop the 1 − *F* (*t*) term in Eq. (S50). However, in this case, the rate of adaptation of the resident ecotype can now be treated as a small correction. On a timescale of ∼1/(*x*_0_(0)− *vT* ^*^)∼1/*s*_*c*_, the invading ecotype will again approach a local equilibrium,

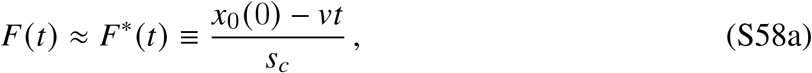

with next-order corrections that are given by

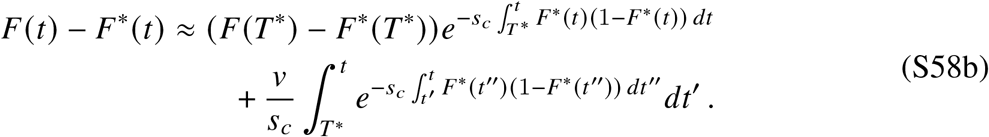

These corrections will be small provided that 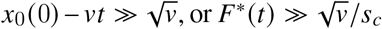, which establishes the region of validity of this fast-relaxation approximation. Note that Eq. (S58) smoothly overlaps with Eq. (S57) in the intermediate region where 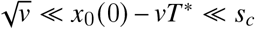, when both approximations are simultaneously valid. For simplicity, we will therefore focus on the small-invader case for the remainder of this section, since the final expressions will be similar.

##### Phase 3: Relaxation to long-term quasi-stable coexistence

At longer times, additional mutations in the invading ecotype will start to become important, and Eq. (S50) will break down. As the founding clone expands, it begins to seed additional fitness classes according to the dynamics described in SI Section 3.3. After each new class establishes, its relative frequency compared to the previous class can be obtained from Eq. (S32), with most of the terms cancelling out:

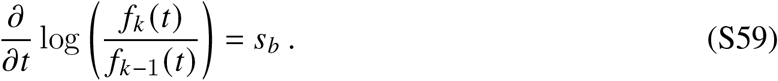

This implies that on a timescale of

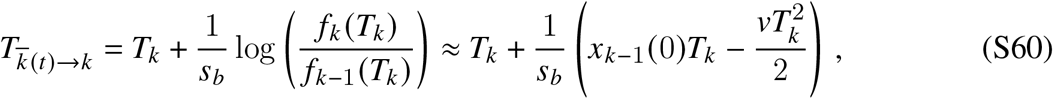

the *k*th class will overtake the *k* − 1th class, and 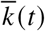 will increase by one. Our analysis of the establishment time distribution in SI Section 4 shows that the second term in this expression is always equal to log (*s*_*b*_ / *U*_*b*_) / *s*_*b*_ *x*_*c*_ *v s*_*b*_ / 2*v*. This yields a simple relation between the establishment time and the switch time,

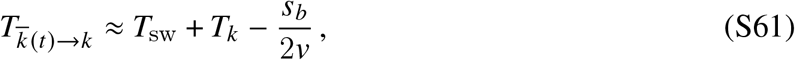

which was referenced in the main text. This result implies that Phase 2 of the trajectory (where 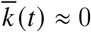) will generally apply up to *t* ≈ *T*_sw_.

###### Ecological impacts of new fitness classes

After this point, the increasing *k t* term requires the full dynamics in Eq. (S49). In the small-invader limit where *F t* 1, this leads to a generalization of Eq. (S56),

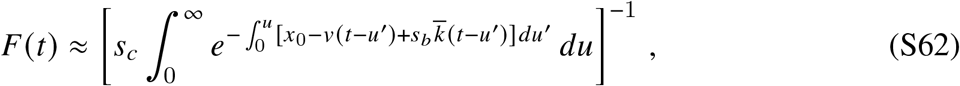

that applies for *t* ≳ *T*_sw_. The dynamics become particularly simple in the case where 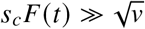, since Eq. (S62) quickly relaxes to the local equilibrium,

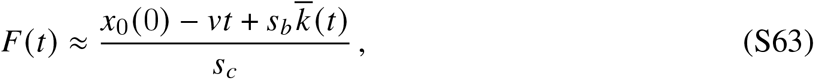

on a timescale much shorter than the rate of change of 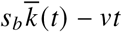. Combining this result with Eq. (S61) leads to Eq. (9) in the main text:

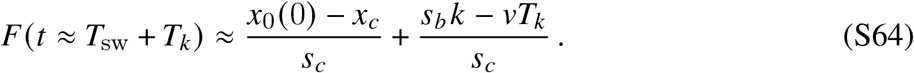

Equations (S62) and (S64) show that the frequency trajectory of the invading ecotype depends on the sequence of establishment times roughly *T*_sw_ generations earlier. For classes with *T*_*k*_ *< T* ^*^, the seeding mutations occurred while the invading ecotype was still at low frequencies, so the establishment times can be predicted from the linear model in SI Section 3.3:

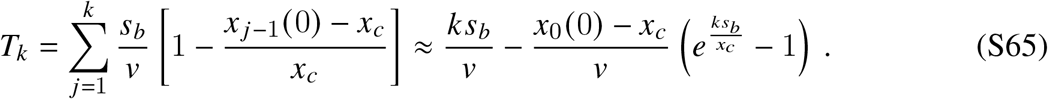

Substituting this result into Eq. (S64), one finds that invading ecotype will start to rebound in frequency after *T*_sw_,

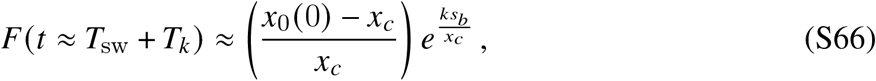

which will continue up to the point where *t* ≈ *T*_sw_ + *T* ^*^. The total number of new classes *k*^*^ that have established by this time can be found by inverting Eq. (S65). When *x*_0_ (0) − *x*_*c*_ ≪ *x*_*c*_, the leading-order contributions are given by

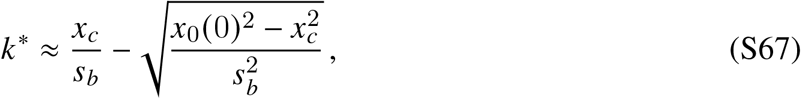

which is close to but strictly smaller than the width of the resident ecotype (*q* ≈ *x*_*c*_/*s*_*b*_). This shows that *F* (*t*) will increase by an 𝒪 (1) factor during the rebound before the ecological feedbacks will start to impact the dynamics of 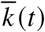.

###### Ecological feedbacks on the establishment process

For fitness classes with *T*_*k*_ *> T* ^*^, the invading ecotype will begin to alter the corresponding establishment times by lowering the effective growth rate of the new mutants. The dynamics are particularly simple in the fast-relaxation case 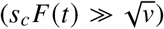, where the invading ecotype remains close to the local equilibrium in Eq. (S62). Substituting this expression into Eq. (S33) yields an effective dynamics for *k* ≥ *k*^*^,

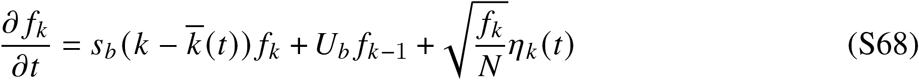

in which the new fitness class primarily competes against the mean fitness of its own ecotype. For *k* = *k*^*^, the corresponding establishment fitness is now smaller than *x*_*c*_,

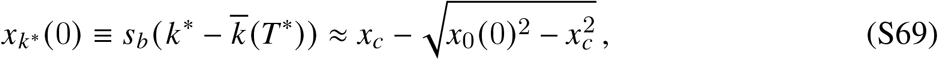

since fewer than *q* ≈ *x*_*c*_ / *s*_*b*_ new classes will have had a chance to establish before *T* ^*^. This implies that the establishment time of the next class will be slightly longer than *s*_*b*_ / *v*, and the corresponding establishment fitness will be slightly lower. This reverses the rebound of the invading ecotype, and causes its frequency to start to decline with time. In principle, these oscillations can continue for several additional cycles until *x*_*k*_ (0) slowly relaxes back to *x*_*c*_. These dynamics are closely related to the fluctuations studied in the non-ecological case in Refs. (52, 54), since they arise from the same dynamical instability between the nose and the bulk of the fitness wave.

###### Connection to the fitness mortgaging picture

We can use these results to estimate the cumulative post-invasion fitness deficit of the invader at time *t* = 2*T*_*sw*_. At this time, the resident ecotype will have advanced in mean fitness by a total amount 2*vT*_*sw*_, while the invading ecotype will be dominated by the class *k*_sw_ with 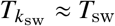. Since the invading ecotype started with a constitutive fitness advantage of 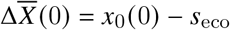 relative to the resident population, its fitness deficit at time *t* = 2*T*_sw_ is given by

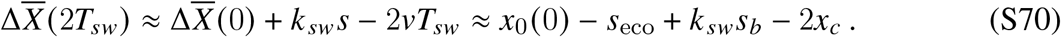

As we have argued, both *x*_0_ (0) and *k*_*sw*_*s*_*b*_ will be close to *x*_*c*_, so the total fitness deficit is approximately 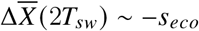. This matches the expectations from the simple “fitness mortaging” argument in the main text. Nonetheless, we have seen that even small values of *x*_0_ (0) − *x*_*c*_ can drive strong transients in the initial phase of the invasion, where the full dynamical picture is required.

### 3.5 Long-term dynamics of established ecotypes

On longer timescales (*t* ≫ *T*_sw_), the cumulative differences in 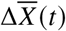 will start to dominate the dynamics in Eq. (S32), and the ecotype frequencies will gradually diverge from their initial post-invasion levels. At this point, it is useful to recast these dynamics as a “race” between two competing fitness waves, which are coupled through the ecological interactions in Eq. (S32). We can formalize this idea by defining the rescaled variables,

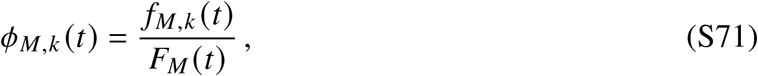

which represent the normalized frequencies of each fitness class within the invading ecotype, along with a corresponding set of variables *ϕ*_*R,k*_ (*t*) representing the resident ecotype’s fitness classes. Substituting this definition into Eq. (S32) leads to a corresponding set of equations for the relative frequencies,

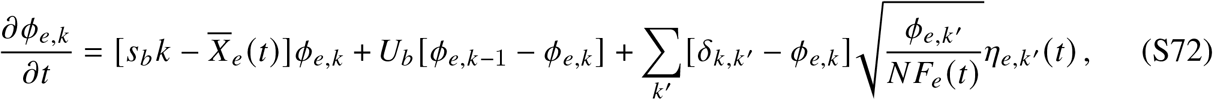

where the interactions between the ecotypes only enter through the time-dependent population size *N*_*e*_ (*t*) ≈ *NF*_*e*_ (*t*). The ecotype frequencies continue to evolve according to Eq. (S49) above,

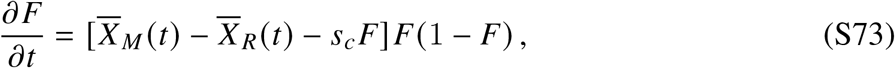

which yields a closed system of equations for predicting 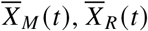, and *F* (*t*).

#### Separation of timescales

Eqs. (S72) and (S73) exhibit a natural separation of timescales when *s*_*c*_*F* (*t*) (1 − *F* (*t*)) 1 *T*_sw_, in which the ecotype frequencies change more slowly than the corresponding dynamics of 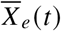. We first make this assumption and solve for the resulting dynamics. We will then use the results to check the regime of validity of the approximation below.

When the ecotype frequencies change sufficiently slowly, Eq. (S72) becomes equivalent to an isolated population with a fixed population size *N*_*e*_ ≈ *NF*_*e*_ (*t*_0_). This suggests that the dynamics of 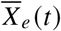 will be well-approximated by existing results from the non-ecological limit in SI Section 3.1. In particular, we we will have 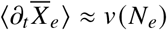 with *v* defined as in Eq. (S29), which implies that the fitness difference 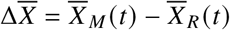 evolves as

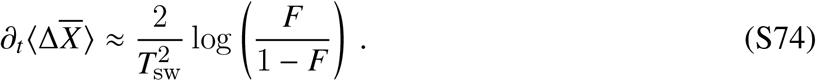

This provides a weak bias in favor of the larger ecotype, which can nevertheless compound on timescales much larger than *T*_sw_. However, as noted in SI Section 3, there will also be fluctuations around this average value. Fluctuations in the establishment of the nose will drive jumps in the mean fitness of each wave ∼*T*_sw_ generations later. Fisher (54) has shown that, on timescales ≫ *T*_sw_, these fluctuations “smooth out” into a Brownian-like noise with a diffusion constant

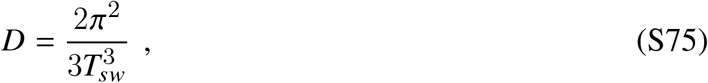

which is approximately independent of *N*. The diffusion constant for the fitness difference 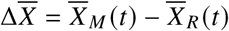 is twice as large, since the fluctuations in each wave contribute independently. This implies that the fitness difference can be described by the stochastic differential equation,

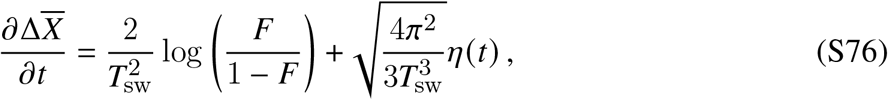

which will apply on timescales that are long compared to *T*_sw_ but short compared to the time required for the ecotype frequencies to change significantly.

We can estimate this latter timescale by returning to Eq. (S73). When *s*_*c*_*F* (*t* (1 *F* (*t*)) 1 *T*_sw_, we can again apply a quasi-static approximation,

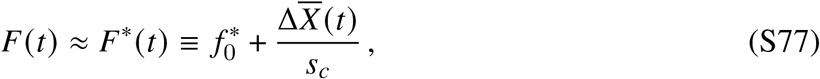

with corrections that we will analyze below. Taking a derivative of this expression and then applying Eq. (S76) yields Eq. (12) in the main text:

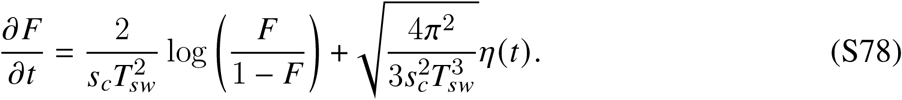

This is similar to the model proposed in Ref. (34), except that it now includes the correct bias term. While the overall magnitude of this bias is small, it is often much larger than the corresponding stochastic term. This has important consequences for the resulting dynamics, which we study in more detail below.

#### Dominant balance analysis of racing fitness waves

We can analyze Eq. (S78) asymptotically in the limit that *s*_*c*_*T*_sw_ ≫ 1. In this regime, most of the non-trivial dynamics occur in a narrow range of frequencies near *F* ≈ 1/2, where the direction of the bias term can switch. Motivated by this observation, we can expand Eq. (S78) around *F* ≈ 1/2 to obtain a linear model for the residual frequency difference *ϵ* (*t*) ≡ *F* (*t*)− 1/2,

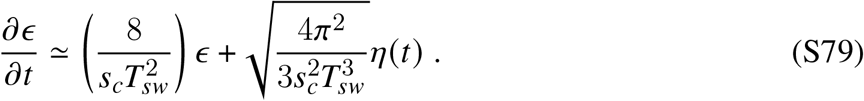

This linearized model has the same functional form as the weak mutation limit in Ref. (34). We therefore quote the relevant results here, and refer the reader to Ref. (34) for more details.

As *F* → 1 / 2, the deterministic difference in the speed of the two ecotypes vanishes, and stochastic fluctuations dominate the dynamics. This suggests that there will be a finite range of frequencies around *F* ≈ 1/2 where we may neglect the effects of the bias term. In this region, wave speed fluctuations over a time interval Δ*t* change the residual difference *ϵ* (*t*) ≡ *F* (*t*)− 1/2 by

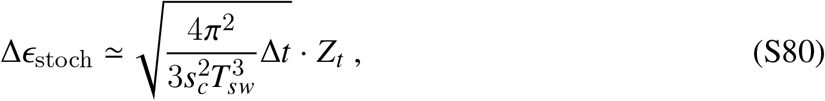

where *Z*_*t*_ is a standard normal variable with (*Z*_*t*_) = 0 and Var(*Z*_*t*_) = 1. Eq. S80 implies that over a timescale 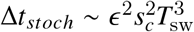, the frequency difference *ϵ* changes by an amount Δ*ϵ*_*stoch*_ ∼ *ϵ*. This approximation is self-consistent provided the deterministic change over the same interval,

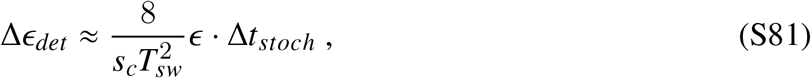

is much smaller in magnitude. Comparing these two terms, we find that the stochastic fluctuations will dominate when |*ϵ* | ≪ *ϵ*_*c*_, where

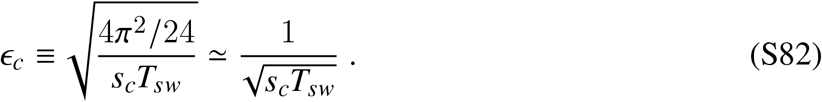

The parameter *ϵ*_*c*_ thus controls the range of the drift-dominated region 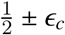 quoted in the main text. This region is much smaller than one when *s*_*c*_*T*_sw_ ≫ 1, justifying our *ϵ* → 0 expansion in Eq. (S79). When *ϵ* ≳ *ϵ*_*c*_, the small differences in the rate of adaptation will start to dominate over fluctuations, causing *ϵ* (*t*) to diverge toward the closest boundary with a characteristic doubling time of 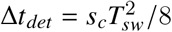. For larger values of *ϵ* (*t*), these dynamics smoothly cross over to the full non-linear bias term in Eq. (S78), which causes the relative frequencies to diverge slightly more rapidly than Eq. (S79). Nevertheless, even when |*ϵ* | ≈ 40%, the higher order terms in a Taylor expansion of Eq. (S78) contribute *<* 40% of the linear term, so this simple approximation holds for a broad range of frequencies.

#### Regime of validity of the quasi-static approximation

Our analysis above shows that *F* (*t*) changes by an O(1) factor on a characteristic timescale of 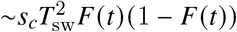. The diffusion approximation in Eq. (S76) will therefore be valid if this timescale is long compared to *T*_sw_, which occurs when *s*_*c*_*F* (*t*) (1 − *F* (*t*)) ≫ 1 / *T*_sw_.

To verify the self-consistency of the quasi-static approximation in Eq. (S77), we can calculate the next-order corrections to *F* (*t*). Writing *δF*(*t*) = *F* (*t*)− *F*^*^(*t*) and expanding Eq. (S73) for small values of *δF*(*t*) yields the linearized equation,

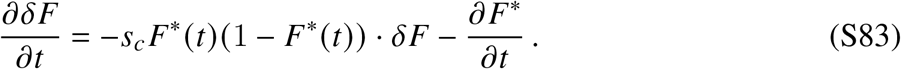

The first term drives *δF* to zero on a characteristic timescale of Δ*t*_*relax*_ ∼ 1 / *s*_*c*_*F*^*^ (*t*) (1 − *F*^*^ (*t*)), which is much shorter than the rate of change of *F*^*^ (*t*) (1 − *F*^*^ (*t*)) when *s*_*c*_*F*^*^ (1 − *F*^*^(*t*)) ≫1 / *T*_sw_. The second term acts like an effective source term, introducing “kicks” away from this local equilibrium. Our analysis above shows that the total magnitude of these kicks over a characteristic relaxation time is bounded by

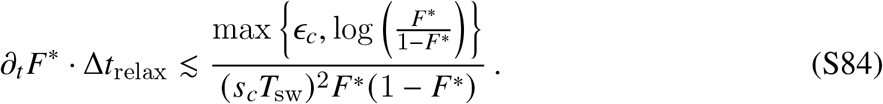

Thus, provided that *s*_*c*_*F*^*^(*t*)(1 − *F*^*^(*t*)) ≫ 1/*T*_sw_, the ecological restoring force will overwhelm the evolutionary destabilization term, and *F* (*t*) will remain close to *F*^*^(*t*).

#### Comparison to the weak mutation limit

The functional form of Eq. (S79) is similar to the weak-mutation limit in Ref. (34),

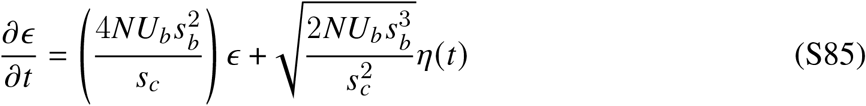

though the underlying parameters are different. In both cases, there is a special range of ecotype frequencies |*F* (*t*) − 1 / 2| *< ϵ*_*c*_ where the dynamics of *F* (*t*) are dominated by random fluctuations in the advance of each wave. In the clonal interference regime, *ϵ*_*c*_ is given by Eq. (S82), which is narrower than the equivalent threshold in SSWM limit (attained by scaling down *N*), by a factor of 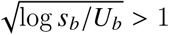, though this factor is typically modest in practice.

More significant are differences in ecosystem lifespan. Measured in “wall-clock” time, coexistence persists for much longer in the weak mutation limit than it does under clonal interference, for the simple reason that the waiting time between successive mutations is asymptotically large. This comparison might suggest that clonal interference weakens long-term coexistence. However, measured in the number of beneficial mutations accumulated in each ecotype prior to collapse, much more “molecular-clock” time passes under clonal interference before the ecosystem fully collapses. For reasonable parameters (*s*_*c*_/*s*_*b*_ ≲ 10 and *Ns* ≳ 10^5^), this number is given by 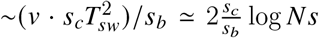, which is at least 10-fold larger than the small-population, weak-mutation limit 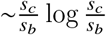. Increased mutational divergence may allow an incipient ecosystem to evolve toward more distinct ecotypes through character displacement, which can reinforce coexistence and delaying or preventing ecosystem collapse. In the presence of diminishing returns epistasis, it can also allow multiple ecotypes to more rapidly approach their fitness optimum, at which point adaptive mutations that would otherwise destabilize coexistence dry up. Consistent with this latter scenario, Ref. (61) showed that larger mutational inputs increased the probability of long-term coexistence in simulations of diversification from an initially maladapted clonal population approaching a fitness peak. These examples show that clonal interference can play a dual role in evolving ecosystems. On one hand, it frustrates the process of initial diversification through the fitness mortgaging effect in SI Section 3.4. However, it can can also promote long-term coexistence between competing ecotypes once they have established.

### 3.6 Recurrent strategy mutations

The previous sections considered the dynamics in the limit that *U*_*α*_ → 0, where at most one new strategy mutation was present in the population at a given time. In this section, we now extend these results to the more realistic case where additional strategy mutations can occur. We start by considering the simplest case, where there are still just two possible resource strategies, *α*_1_ and *α*_2_, and mutations that inter-convert between the two at a total rate *U*_*α*_ ≪ 1. The simulations in Fig. 4 show that, over a broad parameter regime, the clonally interfering ecosystem reaches a quasi-steady ecological state, with high rates of genetic turnover that are driven by periodic cross-invasions between ecotypes. Most of this inter-ecotype genetic turnover is driven by mutations that arise in the leading ecotype [i.e. the one with the higher value of 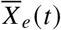] and create new sub-lineages within the lagging ecotype. We now describe a heuristic approach for predicting this ecological steady state.

To fix notation, we let *e* = 1 and *e* = 2 denote the indices of the leading and lagging ecotypes, respectively, and let *F* (*t*) ≡ *F*_2_ (*t*) denote the total frequency of the latter. If the time interval between successful cross-invasions is very long, then we expect that during most of the intervening periods, ecosystem will be well approximated by the quasi-static dynamics in SI Section 3.5. In particular, the balance between the two ecotypes will be set by Eq. (S77), where 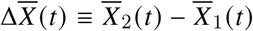 is their current mean fitness difference, and the intra-ecotype frequencies *ϕ*_*e*_ (*X*) will be approximated by a pair of traveling waves with speeds *v*_*e*_ (*t*) = *v* (*NF*_*e*_ (*t*)) and widths *x*_*c,e*_ *t*) = *x*_*c*_(*NF*_*e*_ (*t*)), which depend on the current value of *F* (*t*).

When a new strategy mutation arises in the leading ecotype, it will create a sub-lineage in the lagging ecotype with an initial relative growth rate of

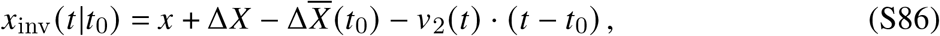

where *t*_0_ is the origination time of the mutation, *x* is the fitness of its genetic background (measured relative to the rest of the leading ecotype), and Δ*X* is the inherent cost or benefit of the mutation. This new sub-lineage will then compete with the lagging ecotype in exactly the same way as a constitutively beneficial mutation with the same initial growth rate. This implies that the establishment probability of the lineage — i.e., the probability that it grows to a large enough size to displace the existing lagging ecotype — is given by the non-ecological formula in Eq. (S25) with *v* = *v*_2_ and *x*_*c*_ = *x*_*c*,2_.

The instantaneous cross-invasion rate is therefore given by

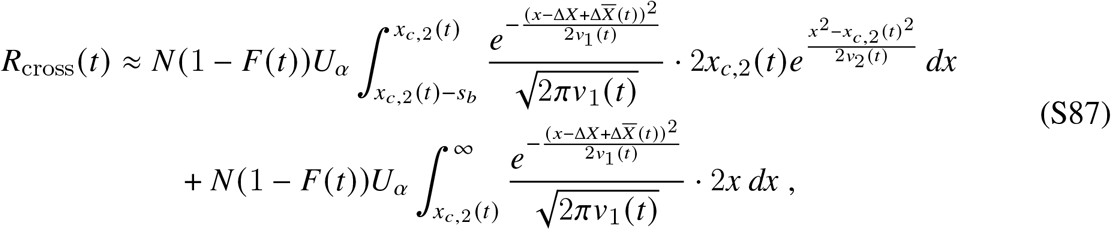

which accounts for the fact that 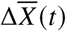 and *F* (*t*) are now slowly varying functions of time. The typical time *T*_cross_ until the first successful cross-invader can then be defined by the heuristic condition in Eq. (14) in the main text:

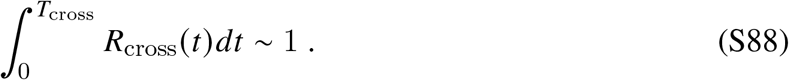

where *t* = 0 denotes the time at which the last successful cross invasion occurred. This heuristic condition defines *T*_cross_ as the time at which the cumulative probability of producing a single strategy mutant destined to cross-invade approaches unity.

To further simplify these expressions, we assume that *s*_*c*_ is sufficiently large that *F* (*t*) remains close to its quasi-steady-state value *F* (*t*) ≈ *F*^*^ (*t*) in the interval between successive cross-invasions, and that this steady-state falls in the deterministic region of Eq. (S76). This implies that the mean fitness difference 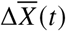 will grow as a linear function of time,

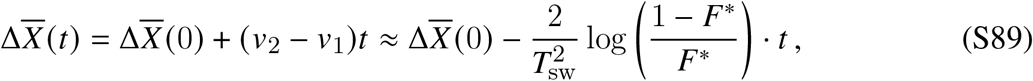

with *v*_1_ and *v*_2_ remaining approximately constant. This linear increase in 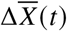 will drive an exponential increase in *R*_cross_(*t*) over the same interval via Eq. (S87). Provided that 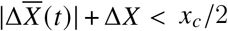, this integral will be dominated by the first term, which reduces to

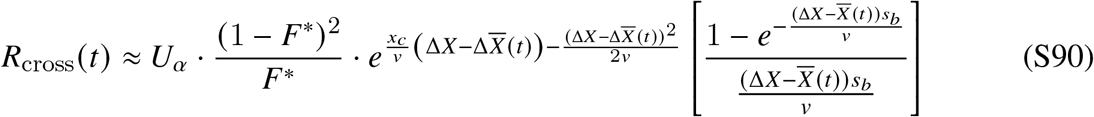

in our parameter regime of interest where *v*_1_ remains close to *v*_2_. This is proportional to the fixation probability of a constitutively beneficial mutation with an effective fitness advantage 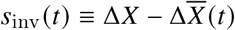.

A self-consistency condition for the ecological steady state can be found by noting that successful cross-invasions must compensate (on average) for the deterministic decline in the mean fitness in Eq. (S89), so that the steady-state values of 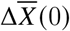 — and therefore *F*^*^ — remain approximately constant over time. The increase in fitness from cross-invasions depends on the establishment fitness *x* of the cross-invader (i.e. where the invader arose relative to the lagging-ecotype’s fitness distribution). As above, provided that the effective fitness advantage of the cross-invader, 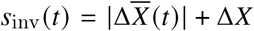, is less than *x*_*c*_, successful cross-invaders will typically arise a distance *v*_2_ / *s*_inv_ *t* below *x*_*c*,2_, and then acquire a subsequent jackpot mutation that pushes their establishment fitness above *x*_*c*,2_. To leading order, the resulting fitness difference *δ* ≡ *x* − *x*_*c*,2_ will have the same distribution as Eq. (S48) in SI Section 3.3, with a typical value of

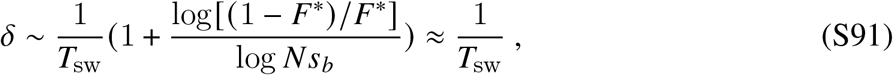

The self-consistency condition 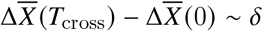 then reduces to

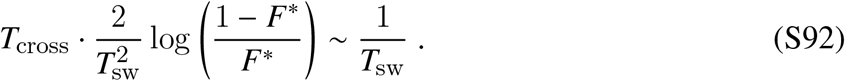

This simplifies to the *T*_cross_ ∼ *T*_sw_ condition quoted in the main text when log (−1 *F*^*^) / *F*^*^ is 𝒪 (1). In practice, we find that a value of *T*_sw_ / 2 provides a reasonable fit to the median value of *T*_cross_ over a wide range of parameters in the simulations in Fig. 4C.

Substituting this expression into Eq. (S88) leads to a transcendental equation for the steady-state value of 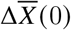 in terms of *U*_*α*_. When log(1 − *F*^*^)/*F*^*^ is 𝒪 (1), the exponential increase of *R*(*t*) during *T*_cross_ amounts to only an 𝒪 (1) factor, so the integral remains of order *R*(0) · *T*_cross_. Solving for 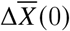 then yields an approximate solution:

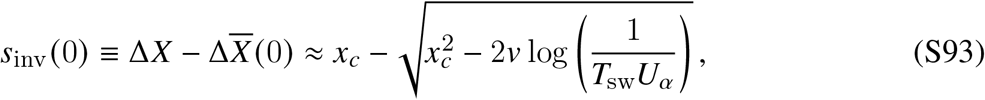

which is self-consistently valid when 1 / *T*_sw_ ≪ *s*_inv_ (0) *< x*_*c*_. When *s*_inv_ ≪ *x*_*c*_, this reduces to Eq. (15) in the main text. The corresponding steady state ecotype frequency is then given by the linear transformation Eq. (S77):

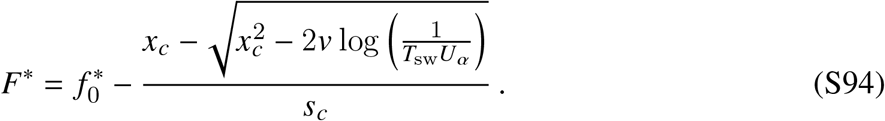

Despite the crudeness of this heuristic argument, we find good quantitative agreement between theory and simulation in Fig. 4C. This agreement also holds for mutations with moderate direct costs (1 / *T*_sw_ |Δ*X*| *s*_*c*_), which are simulated in Fig. S2. Interestingly, the cross-invasion timescale is only marginally affected by a direct cost. While this increases the baseline rates of establishment, our results show that the average fitness difference between ecotypes grows to exactly compensate this cost. Provided that 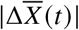 remains smaller than *x*_*c*_, the exponential dependence of *R*_cross_ *t* enforces a similar “typical”, *s*_*c*_-independent cross-invasion timescale as when Δ*X* = 0.

#### Delayed 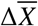 -feedbacks and interfering cross-invaders

This “limit cycle” description of the dynamics, with clearly delineated intervals of *T*_cross_ ∼ *T*_sw_ between successful cross-invaders, omits several complications that we now briefly consider.

Upon first arising near the nose of the lagging ectoype, we have seen that the mutant’s effects on 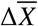 will only manifest after a delay of order *T* ^*^ ∼ *T*_sw_ generations, with *T* ^*^ defined by Eq. (S53). Before this point, its effects are not yet felt by the ecosystem. This implies that an additional mean fitness change due to the difference in population sizes continues to accrue during this time interval. The growing mean fitness difference, in turn, implies that *R*_cross_(*t*) continues to grow exponentially during *T* ^*^, increasing in magnitude by an extra factor of 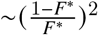 between the beginning and end of the interval. This implies that an additional 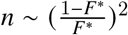 would-be cross-invaders will arise, most of them arising not long before *t* = *T*_cross_ + *T* ^*^, as the first cross-invader reaches macroscopic size.

A majority of these competing cross-invaders will never be detected, since they will clonally interfere with each other. Instead, only the cross-invader with the largest initial jump will ultimately takeover. We can approximate these successful mutants as 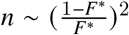 independent draws from the jackpot distribution in Eq. (S48). From extreme value theory, the largest among these will scale as 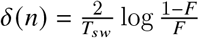, provided *n* is sufficiently small that an exponential approximation of Eq. (S48) is valid out to *δ* (*n*). Thus, after the first cross-invader takes over and increments the mean fitness by ∼1 / *T*_sw_, the largest cross-invader will augment the jump by an additional factor of 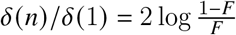 roughly *T*_sw_ generations later. While our minimal argument neglected this additional logarthmic factor, these larger jumps are essential when 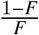 becomes asymptotically large. Considering this log-factor, the deterministic decline in 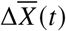 is consistently larger than the jump induced by the first (typical) cross-invader, implying a breakdown of the ecological steady state over repeated cross-invasions. Rather, it is the augmented jump that can ultimately compensate the fitness loss.

These arguments suggest that the simplified “limit-cycle” dynamics are still qualitatively realized, but over several cross-invasions, rather than a single one. These amount to small quantitative corrections for the regimes considered here. Nonetheless, a novel qualitative prediction from this elaboration is that some cross-invasion events should be marked by competition between asymptotically many cross-invaders vying for fixation. We observe signatures of this competition in traces of co-sweeping cross-invaders in Fig. 4B.

#### Breakdown of *T*_cross_ ∼ *T*_sw_ **for large** 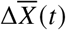

In the main text, we noted that the turnover time begins to rise substantially above *T*_sw_ at small values of *U*_*α*_ (corresponding to a large steady-state value of 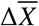). Our self-consistency argument for *T*_cross_ ∼ *T*_sw_ rested on the idea that cross-invaders fix by acquiring a jackpot mutation with *δ* ∼ 1 / *T*_sw_. When 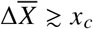, this behavior changes, and the establishment probability will be dominated by the second term in Eq. (S87), with a characteristic fitness increment of 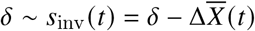. These represents cross-invaders with invasion fitness above the nose of the lagging ecotype. These anomalously fit mutants then races ahead (as in SI Section 3.4) and significantly reduce 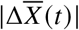. This drives an exponential reduction in *R*_cross_, which must build back up to *R* (*t*) ∼ 1 / *T*_sw_ before another cross-invasion becomes likely. This implies that the the typical lineage survival time begins to increase from ∼1 / *T*_sw_, as observed in the simulations in Fig. 4.

### 3.7 Multi-ecotype dynamics

We can extend the calculation in SI Section 3.6 to settings where multiple types of strategy mutations are available. For simplicity, we will restrict our attention to cases with ℛ = 2 resources, so at most two ecotypes can coexist with each other at a time. As above, we assume that long-term genetic composition of the population will be dominated by one leading ecotype (*e* = 1) and one of several possible lagging ecotypes (*e* = 2,…, *K*). In this case, the key new insight is that the ecological steady-state will no longer be characterized by constant values of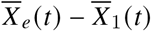 — which will vary depending on which lagging ecotype is currently dominating — but rather by stationary values of the resource availabilities 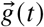 in Eq. (S22b). Under the quasi-steady state in SI Section 3.5, these resource availabilities will satisfy

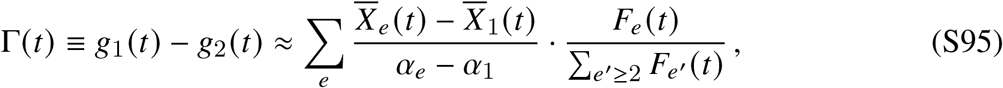

which is typically dominated by the single lagging ecotype with the largest current frequency. The initial growth rate of an arbitrary strategy mutation in such a scenario is then given by a generalization of Eq. (S86),

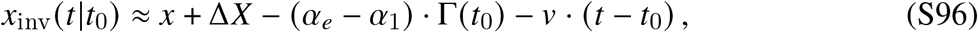

where we have neglected the small differences between the different values of *v*_*e*_ over the lifetime of the new mutation. From this expression, we can identify the ecological fitness advantage in Eq. (16) in the main text.

The rest of the calculation proceeds in a similar manner as SI Section 3.6, leading to the generalized expression in Eq. (17) with *T*_cross_ ∼ *T*_sw_. Thus, even under a continual influx of strategy mutations, the ecosystem tends to organize to a state dominated by ℛ ≈ 2 ecotypes, where there is a small *effective* strategy mutation rate *NU*_*α*,eff_ 1. Analogous to the way that most constitutively beneficial mutations are effectively “wasted” on insufficiently fit genomic backgrounds in traveling fitness waves, most strategy mutations—even those with positive invasion fitness—are wasted in these saturated ecosystems. This sustains longer periods (*>T*_sw_) of strategy stability, with several constitutively beneficial mutations arising within each ecotype before they turn over.

## 4 Solving for the distribution of establishment fitness

In this section, we present the full solution for the distribution of establishment fitnesses in SI Section 3.3. We derive this result in two different ways. We first solve for the distribution formally using moment generating functions and inverse Laplace transforms. We then present a heuristic derivation which captures much of the same behavior. Both approaches heavily draw on previous analyses of the *v* = 0 case (52, 89, 91), with small modifications to incorporate the additional *v* · *t* terms or to show when they are negligible.

### 4.1 Generating function solution

Without loss of generality, we consider the dynamics of a nascent *k* = 1 class,

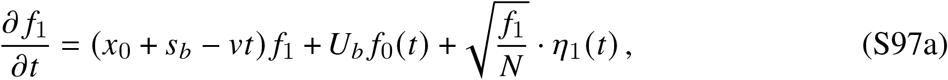

which is fed by mutations from an established *k* = 0 class that is growing deterministically as

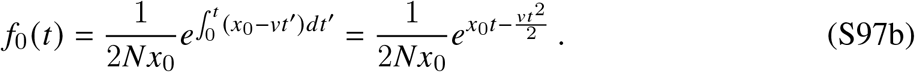

We start by seeking a formal solution for the moment generating function,

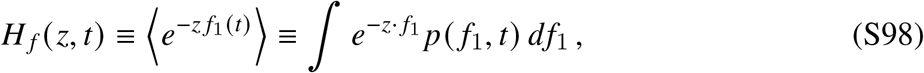

which can be obtained using standard methods. In particular, the Langevin dynamics in Eq. (S97) imply a corresponding PDE for *H* _*f*_ (*z, t*),

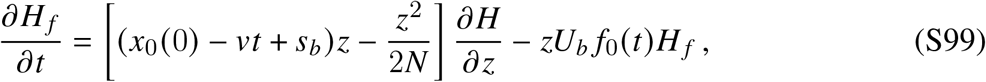

whose general solution is given by

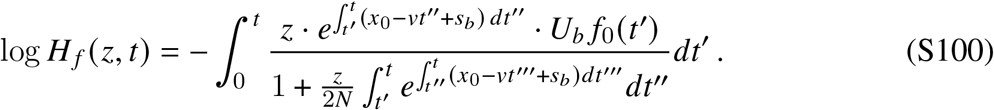

To cast this solution into a more interpretable form, it is convenient to first define the rescaled variable,

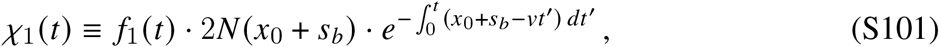

which scales *f*_1_(*t*) by the size of a hypothetical mutation that established at time *t* = 0. After some algebra, the generating function for *χ*_1_(*t*) can be written as

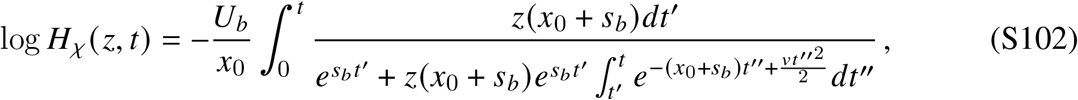

which can be interpreted as a weighted sum over the possible arrival times (*t*,) of all the incoming mutations from *f*_0_ (*t*).

Further simplifications can be obtained in our parameter regime of interest in SI Section 3.3, where 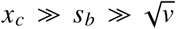 and |*x*_0_ − *x*_*c*_| ≪ *x*_*c*_. In this limit, the two terms in the denominator of Eq. (S102) are rapidly increasing and decreasing functions of *t*,, respectively. This implies that for sufficiently late evaluation times *t*, the integral in Eq. (S102) will be dominated by mutations that arrive near a critical time *t*, = *t*^*^ (*z*), when the two terms are approximately equal to each other. This occurs when

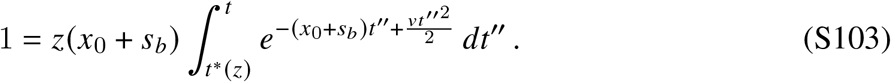

If *t* and *t*^*^ (*z*) are both small compared to *x*_0_ *v*, and *t* ≳ *t*^*^ (*z*) + 𝒪 (1 / *x*_0_), then the integral over *t*^″^ will dominated by its lower limit. Eq. (S103) then reduces to

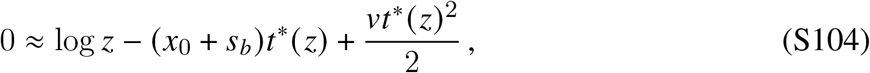

while the integral in Eq. (S102) becomes

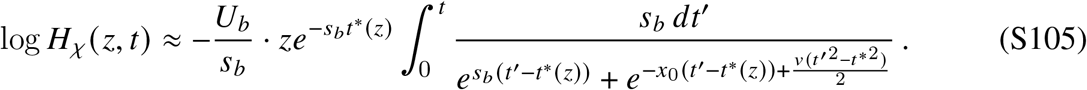

The integrand in Eq. (S106a) decays exponentially away from *t*^*∗*^(𝓏) at rates *s*_*b*_ and *x*_0_ for *t′ > t*^*∗*^(𝓏) and *t′ < t*^*∗*^(𝓏) respectively. Thus, if |*t* − *t*^*∗*^(𝓏)| ≳1/*s*_*b*_ ≫ 1/*x*_0_, the integral becomes dominated by the shoulder above *t*^*∗*^(𝓏) and rapidly converges to one, leaving

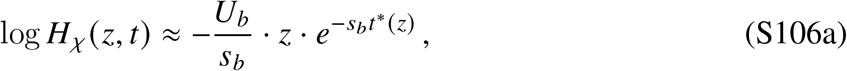

with *t*^*∗*^(𝓏) defined by

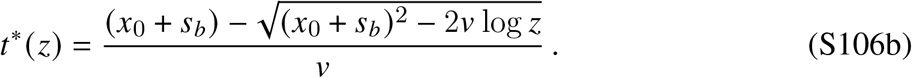

#### Inverting the generating function

The probability density of 𝒳_1_ can be obtained from the moment generating function via an inverse Laplace transform,

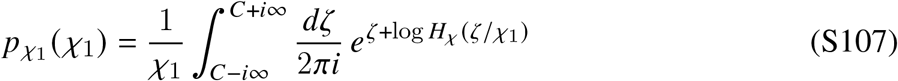

If we assume that the integral is dominated by values of ζ with log ζ ≪(*x*_0_ + *s*_*b*_) ^2^ /*v* + log 𝒳, we can Taylor expand the square root in Eq. (S106b) to obtain

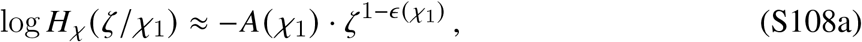

where we have defined the 𝒳_1_-dependent quantities

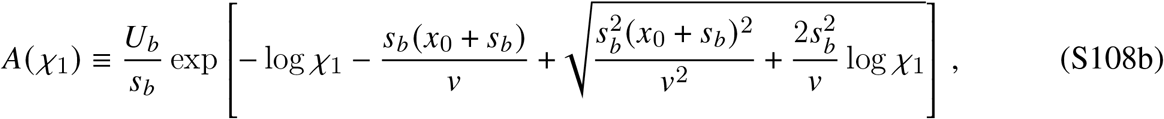

and

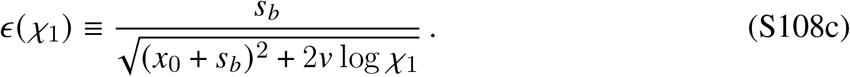

This is similar to the Levy-like distribution in the *v* = 0 case in Ref. (52), except that the *ϵ* parameter now depends on 𝒳_1_.

The inverse Laplace transform can be evaluated by setting the branch cuts of this function – infinitely many for irrational values of *ϵ* (𝒳_1_) – to the negative real line, (− ∞, 0], and closing the contour while remaining on the principal branch Arg ζ ∈ (−*π, π*): starting at +*i*∞ and moving counterclockwise to ™ ∞, coming back to the origin an *ϵ* above the negative real axis, wrapping around the origin and back out below the negative real axis, and closing with a quarter-circle from ™ ∞ to ™*i*∞. Since this closed Hankel-type contour contains no poles, its integral evaluates to zero. The only non-zero contributions come from Eq. (S107) and the paths from (™ ∞ +*iϵ*, +*iϵ*) and ™*iϵ, ™* ∞ − *iϵ*) above and below the negative real axis, respectively. Writing ζ = *ye*^±*iπ*^ depending on whether we are above or below the negative real axis, we find that

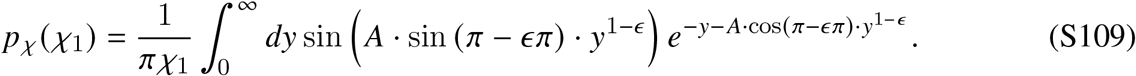

We can use this intermediate result to solve for the corresponding distributions of the establishment time (*T*_1_) and the establishment fitness (*x*_1_).

**Distributions of** *T*_1_ **and** *x*_1_. The definition of the establishment time in Eq. (S40) yields a relation between 𝒳_1_ and *T*_1_:

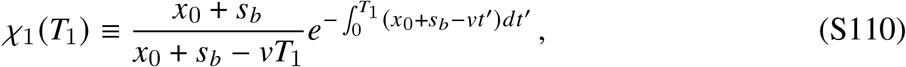

which can be combined with the definition of the establishment fitness,

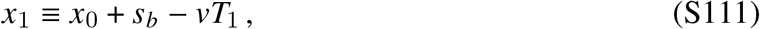

to obtain a corresponding relation between 𝒳_1_ and *x*_1_:

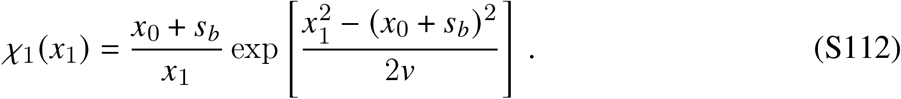

The distribution of *x*_1_ can then be obtained from Eq. (S109) using the change of variables theorem. In our parameter regime of interest where |*x*_1_ ™ *x*_*c*_| ≪ *x*_*c*_, the leading order contributions can be expressed in the compact form,

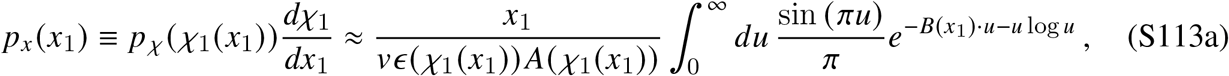

where the function *B*(*x*_1_) is defined by

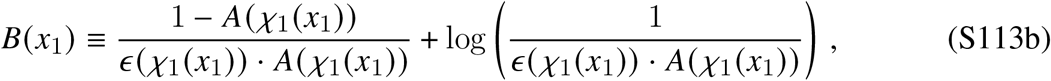

and the *A* and *ϵ* functions reduce to

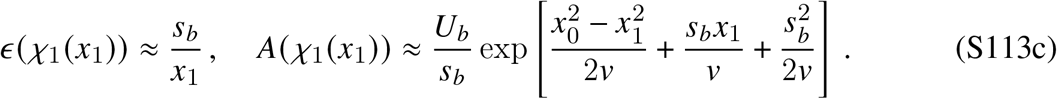

An analogous distribution for the establishment time *T*_1_ can be obtained from the linear transformation in Eq. (S111).

#### Typical values

The integral in Eq. (S113) attains its maximum value when *B*(*x*_1_) = *B*^*∗*^ ≈ −0.2, and falls off by an order of magnitude when |*B*| ≳ 𝒪 (1). This implies that the distribution of *p*_*x*_ (*x*_1_) will dominated by values of *x*_1_ where *A* is close to one:

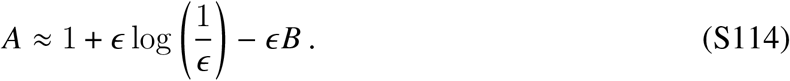

Letting 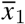 denote the special point where 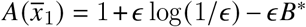, one can expand the resulting expressions to show that

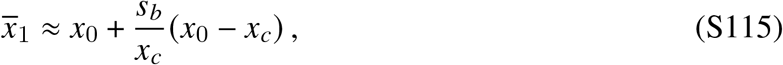

with corrections that are smaller by additional factors of *s*_*b*_/*x*_*c*_ ≪ 1. The typical spread around this value occurs for 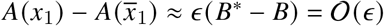, which can be similarly expanded to obtain

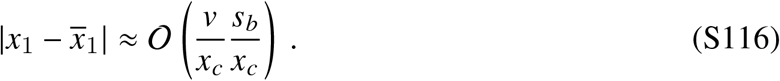

The linear relation in Eq. (S111) then yields an analogous expression for the typical establishment time:

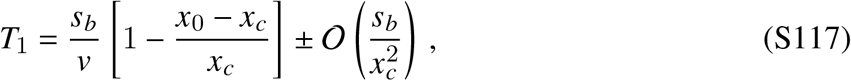

which we quoted in SI Section 3.3.

#### Rare fluctuations

The probability of a larger negative fluctuation in *x*_1_ (corresponding to an anomalously late establishment time) decays extremely rapidly with 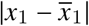, since sine term in Eq. (S113) leads to positive and negative cancellations for large negative *B*. However, this integral decays much more slowly for large positive *B*, which occurs when 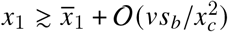. In this case, a Laplace approximation to Eq. (S113) yields

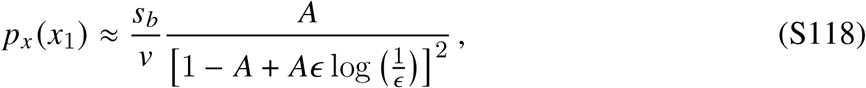

This further reduces to the jackpot distribution in Eq. (S46) in SI Section 3.3 when *A*(*x*_1_) ≪ 1.

### 4.2 Heuristic version

We can also derive many of these same results using a heuristic approach, similar to Refs. (52, 91), which emphasizes the arrival times of the individual mutations that contribute to the next fitness class. Each successful mutation founds a clonal sub-population that grows as

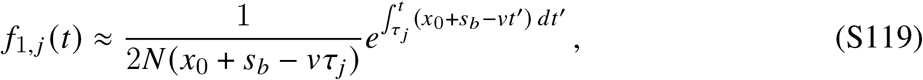

where *τ*_*j*_ is the corresponding establishment time of the *j* th such mutation. Since these individual clones must all sum up to *f*_1_ (*t*), we can obtain a corresponding relationship between the establishment times of the individual mutations and the establishment time of the entire fitness class,

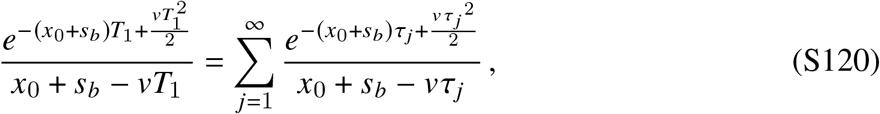

where we have divided out by the common factor of 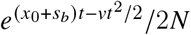.

#### Typical values

The typical values of *T*_1_ will correspond to cases where the sum in Eq. (S120) is dominated by contributions from a large number of mutations. This allows us to approximate the sum by an integral,

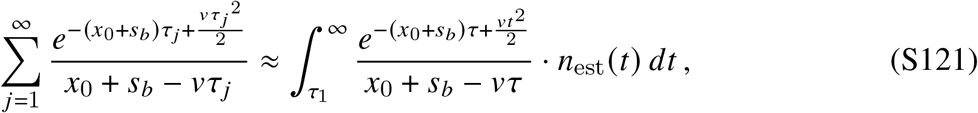

where *n*_est_ (*t*) = *NU*_*b*_ *f*_0_ (*t*) · 2 (*x*_0_ + *s*_*b*_ – *vt*) is the average density of (successful) mutations that are produced at time *t*. Substituting this function into Eq. (S121), many of the time-dependent factors cancel out, and the integral reduces to the simpler form,

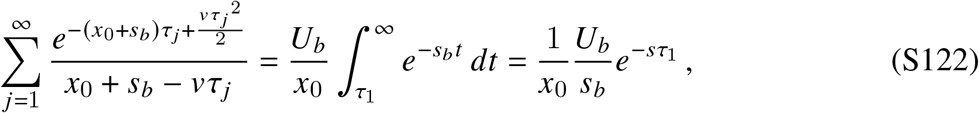

which is dominated by mutations that arise within ∼1/*s*_*b*_ generations of *τ*_1_. The inclusion of this lower bound is critical: since *n*_est_ (*t*) is a rapidly increasing function of *t*, many early times will correspond to cases where we are unlikely to sample any mutations.

We can estimate the typical value of *τ*_1_ from the heuristic condition that at most one mutation will establish before this point

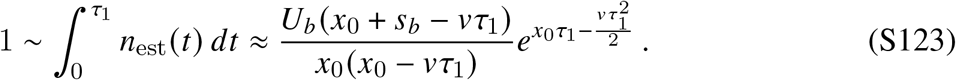

Combining this result with Eqs. (S120) and (S122), we find that the typical value of *T*_1_ is given by

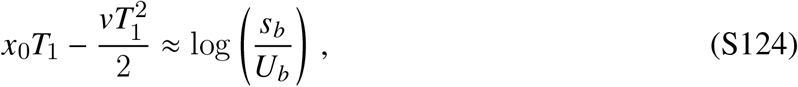

with corrections that are much less than one in the limit that 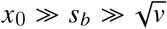. The establishment fitness *x*_1_ ≡ *x*_0_ + *s*_*b*_ − *vT*_1_ then satisfies a similar condition,

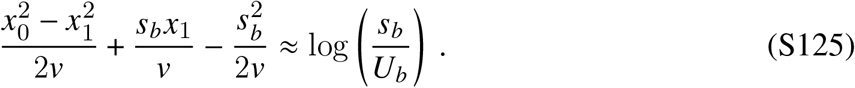

The critical point *x*_*c*_ corresponds to the location where *x*_1_ = *x*_0_, which occurs when

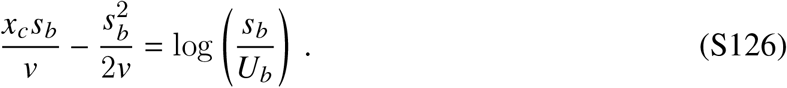

This provides an alternative derivation of Eq. (S26) in SI Section 3.1. We can use this definition of *x*_*c*_ to rewrite Eqs. (S124) and (S125) as

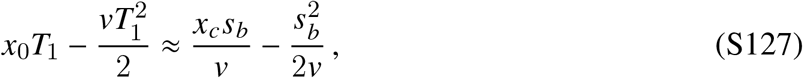

and

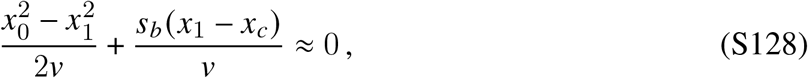

respectively, which reduce to Eqs. (S43) and (S44) in SI Section 3.3 in the limit that |*x*_0_ − *x*_*c*_ | ≪ *x*_*c*_.

#### Rare fluctuations

Since the typical values of *T*_1_ and *x*_1_ are dominated by many independent mutations, the corresponding fluctuations in these quantities will also typically be small. However, in rare cases, the first mutation will arise sufficiently early that it will dominate the sum in Eq. (S120), so that *T*_1_ ≈ *τ*_1_. The probability density of these events is simply *n*_est_ (*T*_1_), which reduces to the jackpot distribution in Eq. (S46) in SI Section 3.3 after changing variables from *T*_1_ to *x*_1_:

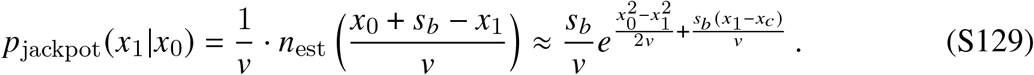

## 5 Simulations and numerical methods

We simulated Eq. (S8) in a Wright-Fisher-like framework, similar to the approach described in Ref. (34). Each simulation begins with a clonal population of size *N* in an environment with fixed resource supply vector 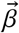. Each subsequent generation is composed of two steps:

1. **Selection step:** For each strain *μ*, the initial number of individuals in the next generation is sampled from a Poisson distribution with mean

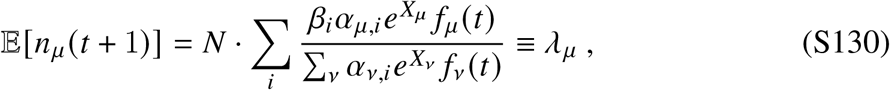

where *f*_*μ*_ (*t*) ≡ *n*_*μ*_ (*t*)/Σ_*ν*_ *n*_*ν*_ (*t*) denotes the relative frequency of the strain in the current generation. Note that the expected value of the sum is E[Σ_*μ*_ *λ*_*μ*_] = *N*, ensuring that the fixed population size assumption is respected on longer timescales up to 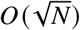 fluctuations.
2. **Mutation step:** After the reproduction step, each individual in the population acquires a mutation with probability *U* = *U*_*b*_ + *U*_*α*_. A fraction *U*_*b*_/*U* of these mutations are constitutively beneficial mutations and produce a new strain with phenotype 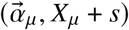, with *s* drawn from the distribution of fitness effects *ρ*_*b*_ (*s*). For most of this work, we consider the single-s distribution *ρ*_*b*_ (*s*) = *δ*(*s* − *s*_*b*_); extensions to exponentially distributed fitness effects are considered in Fig. S3. The remaining fraction of mutations (*U*_*α*_/*U*) are resource strategy mutations, and produce a new strain with phenotype 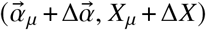, with 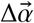 drawn from the resource strategy mutation kernel 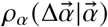. In all simulations, we considered a discrete set of *L* strategies 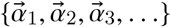, such that the strategy mutation kernel was specified by an *L*×*L* matrix.

To simulate the asymptotically large population sizes in Fig. 3), we ignored birth-death stochasticity in subpopulations with *λ*_*μ*_ *> e*^20^, and set their updated population sizes to be equal to *λ*_*μ*_. Similarly, for subpopulations where the expected number of mutants is larger than *e*^20^, we set the number of new mutants equal to the deterministic expectation.

In most simulations, *μ* indexes “phenotype classes”, i.e. all cells sharing the same fitness *X*_*μ*_ and resource uptake strategy 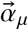, regardless of their evolutionary relatedness. When fitness effects and strategy changes are drawn from discrete distributions (as is the typical case in this study), coarse-graining over distinct lineages sharing the same fitness and strategy dramatically speeds up simulations. However, *μ* may also index individual lineages, each composed of cells with identical genotypes deriving from the same ancestor. We utilize lineage-based simulations to produce the Muller plots in Fig. 1. Additionally, we used a hybrid phenotype/lineage tracking approach to measure inter- and intra-lineage turnover dynamics in Figs. 4 and 5, which is described in Section 5.2 below.

### 5.1 Simulating rare invasions

We used this simulation scheme to study the dynamics and establishment probability of a new strategy mutation in an initially mono-ecotype population (Figs. 2 and 3). In all simulations, we considered a strategy mutation as “established” in the population if it reached a threshold population-wide frequency *F*_est_, which was typically set to *F*_est_ = 10%. This is an operational definition: it is large enough that the mutant lineage has established its own traveling wave (since *NF*_est_*U*_*b*_ ≫ 1), but still small compared to the expected ecological equilibrium *f* ^*∗*^ in the absence of clonal interference.

As noted above, each simulated population was initialized with a clone of a single strategy and fitness. To avoid considering invasions during the transient dynamics associated with the build-up of the traveling wave (52), we allowed the mono-strategy population to evolve by fitness mutations alone for *T*_burn_ = 10 · *T*_sw_ generations, with *U*_*α*_ set to 0. After the burn in, *U*_*α*_ was set to a finite but sufficiently small rate. In the limit that *U*_*α*_ → 0, the establishment probability *p*_est_ can be calculated by the relation

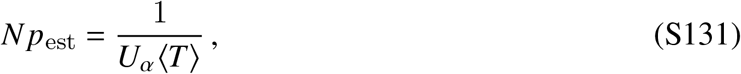

where ⟨*T*⟩ is the average time to produce a strategy mutant destined to establish, with the average taken over all possible realizations of the post-burn-in traveling wave. This correspondence is strictly exact only as long as at most one strategy mutation is segregating in the population at any one time; otherwise, two or more strategy mutations (weakly) bias each other’s competition against the resident. However, it is still sufficiently accurate so long as the collective frequency of new strategy mutations remains small over the course of simulation, up to the point when the establishing lineage reaches establishment size. In practice, this is obtained for sufficiently small values of *U*_*α*_ that the typical time between burn-in and establishment is much larger than *T*_sw_: this reflects the fact that mutations that are not destined to establish will typically go extinct within *T*_sw_ generations.

To ensure that this condition was satisfied, we simulated a series of establishment runs under a sequence of strategy mutation rates 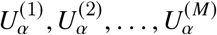, producing corresponding sequence establishment times *T* ^(1)^, *T*^(2)^,…, *T* ^(*M*)^. After each establishment run *i*, we computed a rescaled strategy mutation rate

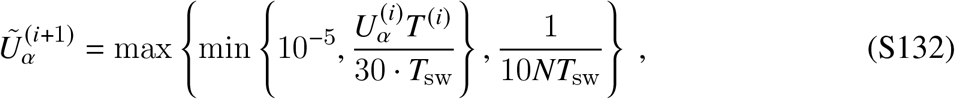

which rescales *U*_*α*_ to target a desired establishment time of 30 · *T*_sw_. The maximum rescaled rate 10^−5^ ensures that *F* (*t*) remains small up to the time of establishment, while the minimum rescaled rate 1 10*NT*_sw_ prevents excessive waiting between mutation attempts and arithmetic underflow when *s*_inv_ ≳ *x*_*c*_. The strategy mutation rate of the next simulation run was then set to the median value of 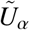 of the 10 previous runs:

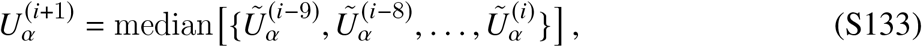

with 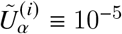. We allowed the realized strategy mutation rates to equilibrate over 50 initial simulations, and then estimated *p*_est_ from the average of all subsequent runs:

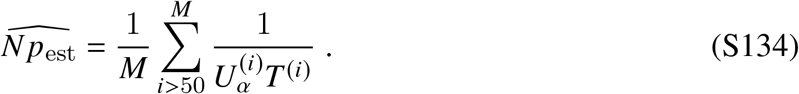

### 5.2 Cross-invasion dynamics during ecotype coexistence

To simulate the dynamics of coexistence and cross-invasion in Figs. 4 and 5, we used a pseudo-lineage tracking approach that was amenable to simulation over a discrete strategy landscape in the high strategy mutation rate regime. In this approach, the subpopulation in each ecotype 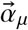 is distributed over *k* = 50 dummy lineages (or “rows”). Between cross-invasion events, the population density of a given strategy will be concentrated in just a handful of rows, corresponding to previously successful cross-invaders that are continuing to accrue beneficial mutations. Each generation, new strategy mutations from other ecotypes are randomly distributed across the *k* ≫ 1 rows of the target ecotype. When a new strategy mutation establishes and replaces an existing lineage, this strategy mutation will typically arise in one of the remaining rows that did not contain the originally dominant population. This provides an efficient means to track successful cross-invasions, without the computational complexity of tracking every individual lineage.

To count successful cross-invasions, we initialized a monoclonal population and allowed a burn-in for 10 · *T*_sw_ generations. After burn-in we indexed the dominant rows in each strategy. Over subsequent generations, we denoted a successful strategy cross-invasion when the population density in one dummy row of a strategy (i) exceeded that of the current dominant row and (ii) was itself *>* 0.1%. In Fig. 5, we also tracked the dominant strategies in the population at a given time: the ecosystem was defined as in the 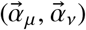 state when the total frequency of individuals with 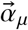 or 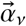 exceeded the frequency of all other ecotypes by at least 10-fold.

### 5.3 Numerical methods

To compare our theoretical predictions with simulations, we first used Eqs. (S29) and (S26) to calculate *v* and *x*_*c*_ from the underlying values of *N, U*_*b*_ and *s*_*b*_. We also calculated the corresponding sweep timescale *T*_sw_ ≡ *x*_*c*_ */ v*, which is approximately *N ™*independent. We used these pre-calculated values to estimate the relevant fitness- and time-scales denoted in Figs. 2-5 in the main text. Likewise, the theory curves in Fig. 4D were obtained by substituting our estimates for *x*_*c*_, *v*, and *T*_sw_ into Eq. (S94). For the inset of Fig. 5C, the mutation rates leading to equal amounts of time spent in the (*α*_1_, *α*_2_) and (*α*_1_, *α*_3_) states (Φ = 1 / 2) were estimated from a cubic spline interpolation of the simulated Φ values in Fig. 5C.

